# Nrf2 modulates the hybrid epithelial/mesenchymal phenotype and Notch signaling during collective cancer migration

**DOI:** 10.1101/2021.04.21.440858

**Authors:** Samuel A Vilchez Mercedes, Federico Bocci, Ninghao Zhu, Herbert Levine, José N Onuchic, Mohit Kumar Jolly, Pak Kin Wong

## Abstract

Hybrid epithelial/mesenchymal cells (E/M) are key players in aggressive cancer metastasis. It remains a challenge to understand how these cell states, which are mostly non-existent in healthy tissue, become stable phenotypes participating in collective cancer migration. The transcription factor Nrf2, which is associated with tumor progression and resistance to therapy, appears to be central to this process. Here, using a combination of single cell biosensors, immunocytochemistry, and computational modeling, we show that Nrf2 functions as a phenotypic stability factor for hybrid E/M cells by inhibiting a complete epithelial-mesenchymal transition (EMT) during collective cancer migration. We also demonstrate that Nrf2 and EMT signaling are spatially coordinated near the migrating front. In particular, computational analysis of an Nrf2-EMT-Notch network and experimental modulation of Nrf2 by pharmacological treatment or CRISPR/Cas9 gene editing reveal that Nrf2 stabilizes a hybrid E/M phenotype which is maximally observed in the interior region immediately behind the leading edge. We further demonstrate that the Nrf2-EMT-Notch network enhances Dll4 and Jagged1 expression near the leading edge, which correlates with the formation of leader cells and protruding tips. Altogether, our results provide direct evidence that Nrf2 acts as a phenotypic stability factor in restricting complete EMT and plays an important role in coordinating collective cancer migration.

## 1. Introduction

A most devastating feature of cancer is its ability to migrate and invade adjacent tissues [1]. During invasion by carcinomas, cancer cells can undergo an epithelial-mesenchymal transition (EMT) to gain mesenchymal traits, such as increased motility and invasiveness [2]. Emerging evidence reveals that EMT is not an irreversible, binary process; in contrast, EMT is a reversible transition process with one or multiple hybrid, or partial, epithelial/mesenchymal (E/M) states which can help coordinate the collective invasion of cancer cells [2,3]. These intermediate states arise due to the complex dynamics of cell fate circuits encompassing mutually inhibiting microRNA and EMT transcription factors [4,5]. Hybrid E/M phenotypes have been associated with more stem cell-like traits, which include resistance to treatment and enhanced aggressiveness in comparison with purely mesenchymal or epithelial phenotypes [6,7]. The clinical significance of the hybrid E/M phenotype in collective cancer invasion is evidenced by analyses of circulating tumor cell clusters exhibiting both mesenchymal and epithelial phenotypes [8,9]. Furthermore, the hybrid E/M phenotype has been associated with increased metastatic potential and poor clinical outcomes [10,11].

EMT is a complex process involving various signaling pathways [2,3]. Recent mathematical modeling and experimental analyses have demonstrated that a set of phenotypic stability factors (PSFs) can promote and stabilize hybrid E/M state(s) [12,13]. For instance, the transcription factor nuclear factor erythroid 2-related factor 2 (NFE2L2), commonly referred to as Nrf2, is implicated in EMT regulation [14] and is associated with poor clinical outcomes in cancer patients [15,16]. By integrating computational and experimental approaches, we have previously shown that Nrf2 downregulation destabilizes the hybrid E/M state and prevents collective migration in multiple cancer cell lines, while Nrf2 expression stabilizes a hybrid E/M phenotype that co-expresses epithelial and mesenchymal markers [17].

Notch signaling has also been separately implicated in the regulation of EMT [13,18] and the formation of leader cells during collective cancer invasion [19]. Notch is an evolutionarily conserved mechanism, which regulates cell-fate differentiation and cell-cell coordination [20]. When activated in cancer cells, members of the Notch family, such as Notch1 and its ligands Dll4 and Jagged1, are linked to proliferation, survival, and progression [21]. For instance, Dll4 mRNA is upregulated in leader cells, while Notch1 is upregulated in follower cells [22–26]. Moreover, despite Dll4 and Jagged1 having opposite functions in the regulation of angiogenesis [27], Jagged1 promotes MYO10 driven filopodial persistence and invadopodium formation in leader cells [28]. Computational models of Notch1-Jagged1 signaling have predicted the formation of collectively migrating clusters with hybrid E/M phenotypes [29]. Finally, Notch signaling is directly coupled to Nrf2 expression via reciprocal positive feedback [30,31]. The complex interplays between Nrf2, EMT, and Notch signaling and their roles in collective cancer migration remain poorly understood.

In this study, we investigate the influence of Nrf2 on cancer cells during collective migration, using a combined experimental-computational approach. We experimentally characterized Nrf2 and the EMT related markers, E-cadherin and ZEB1, during collective migration of urothelial bladder cancer cells with immunocytochemistry, fluorescence in situ hybridization (FISH), and a double-stranded single cell biosensor [32], The expression of Nrf2 was modulated by a potent Nrf2 activator, sulforaphane (SFN), or by CRISPR-Cas9 gene editing. The influence of Nrf2 on EMT and Notch signaling was also investigated by a computational model of the Nrf2-EMT-Notch signaling circuit. Specifically, we coupled the intracellular Nrf2-EMT circuitry with the Notch cell-cell communication pathway, which consists of the Notch transmembrane receptor, the Notch intracellular domain (NICD), and the Notch ligands Dll4 and Jagged1. Computations were carried out in a multicellular lattice model that captures the main geometrical features of wound-induced collective cell migration. Members of the Notch family, including Notch1, Jagged1, and Dll4, were measured at the protein and/or mRNA level in single cells. Lastly, we measured phenotypic behaviors, including the formation of leader cells, the morphology of the migrating edge, and the migration speed, in relationship to Nrf2 modulation. The results reveal the important role of Nrf2 in coordinating the hybrid E/M phenotype in cancer cells during collective migration.

## 2. Materials and Methods

### 2.1 Cell culture and reagents

The human RT4 cells (labeled as WT) were purchased from American Type Culture Collection (ATCC, USA). The CRISPR-Cas9 knockout cell pool RT4-Nrf2-KO (labeled as KO) was obtained from Synthego, CA. DL-sulforaphane (cat. #s4441, Sigma Aldrich, USA) was dissolved in DMSO (cat. #D8418, Sigma Aldrich, USA) according to manufacturer’s instructions. DL-sulforaphane was added to the RT4 cells (labeled as SFN) at a final concentration of 7.5 μM immediately after the wound scratch assay. All cells were cultured in McCoy’s 5A medium containing 10% FBS and 0.1% Gentamicin (Fisher Scientific, Hampton, NH). Cells were maintained at 37°C in 5% CO_2_, and media were refreshed every 2 days. The following gRNA targeting exon 2 was used for NFE2L2-KO: AUUUGAUUGACAUACUUUGG. Knockout cells showed a predicted functional knockout of 63% which was confirmed by Synthego through RT-qPCR showing 75% editing efficiency post expansion at passage 4. All experiments were done between passages 5–8 for the CRISPR/Cas9 Nrf2-KO pool cells in order to maximize the population of knockout cells within the pool. All experiments were performed in polystyrene 24-well plates (cat. # 07-200-740, Fisher Scientific, Hampton, NH).

Double-stranded locked nucleic acid (dsLNA) biosensors and synthetic targets for calibration were synthesized by Integrated DNA Technologies (San Diego, CA). The sequences are available in Supplementary Table 1. The following reagents were used to perform FISH: Stellaris RNA FISH Hybridization Buffer (cat. #SMF-HB1-10), Stellaris RNA FISH Wash Buffer A (cat. #SMF-WA1-60), and Stellaris RNA FISH Wash Buffer B (cat. #SMF-WB1-20). All FISH reagents were acquired from Biosearch Technologies. Transfection reagents for the dsLNA biosensors were acquired from Thermofisher scientific.

### 2.2 Immunocytochemistry

Cells were washed with warmed 1x Phosphate-Buffered Saline (PBS) twice, followed by fixation with chilled 4% paraformaldehyde (Sigma) in PBS for 15 minutes. All reagents were kept cold past this point, and incubation was performed at room temperature. Cell permeabilization was performed with 1% Triton X-100 in PBS for 10 minutes followed by a blocking step with 3% bovine serum albumin (BSA) in PBS for 30 minutes. The cells were incubated overnight at 4°C with the primary antibodies and then incubated in the dark for 2 hours against the secondary antibodies. Primary antibodies used were Nrf2 (1:100, cat. #AF3925, R&D Biosystems), E-cadherin (1:50, cat. #M3612, Agilent Dako), ZEB1 (1:100, cat. #ab124512, Abcam), Jagged1 (1:50, cat. #sc-390177, Santa Cruz Biotechnology), Notch1 (1:100, cat. #ab8925, Abcam), Dll4 (1:100, cat. #PA585931, Thermofisher Scientific). Secondary antibodies used were Alexa-fluor conjugated secondary antibodies (1:1000, Life technologies). The antibodies were all diluted in 3% BSA solution. Wells were washed 3 times with PBS in between each step. Cells were examined using a laser scanning confocal microscope (Leica TCS SP8; Leica Microsystems, Wetzlar, Germany) immediately after the last washing step.

### 2.3 Wound-Healing Assay

The scratch wound healing assay was performed to determine cell migration using confluent cultures. Briefly, at 100% confluency, the monolayer was scratched with a sterile 200 μL pipet tip, and the media were refreshed for all wells. Cells were washed with warm 1x PBS before and after wounding. Images were acquired at 0, 24 and 48 hours. The experiments were repeated at least 3 times, and the mean and standard deviation were calculated using ImageJ. Cell migration was expressed as the migration rate in microns per hour (μm/h): (original scratch width – final scratch width)/time.

### 2.4 Single cell gene expression analysis and transfection

The dsLNA biosensors were used to measure mRNA and microRNA (miRNA) expression profiles of target genes near the migrating front of the monolayer (Supplementary Figure S1). The design, characterization, and protocol were described previously [24,32]. Briefly, the complementary sequence to the target mRNA/miRNA (labeled as probe) is labeled with a fluorophore at the 5’ end. A complementary sequence with a quencher at the 3’ end (labeled as quencher) is designed. All sequences were verified through the NCBI Basic Local Alignment Search Tool for nucleotides (BLASTn). A random probe with no known intracellular targets was also developed as a negative control. For transfection, the probe and quencher were dissolved in 10 mM Tris-EDTA buffer and 0.2 M NaCl before mixing at a 1:2.5 ratio. Then, the probe and Lipofectamine RNAiMAX reagent (cat. #13778075, Thermofisher) were diluted in Opti-MEM media (cat. #31985062, Thermofisher) according to the manufacturer’s protocol. Cells were seeded in a 24-well plate and transfected once they reached 90-95% confluency. Each well contained a total of 1 μg probe with 2 μL Lipofectamine RNAiMAX. The dsLNA biosensors targeting different genes were transfected in separate wells.

### 2.5 Fluorescence in situ hybridization

FISH was used to measure mRNA and miRNA expression of target genes in fixed cells near the wound boundary. The FISH assay was performed according to manufacturer’s instructions with the probes designed for the single cell biosensors. Briefly, 24 hours after the wound scratch assay, the cells were fixed using 3.7% formaldehyde in 1x PBS and incubated at room temperature for 10 minutes. Cells were then washed twice with 1x PBS and permeabilized using 70% ethanol in deionized (DI) water for at least 1 hour at 4°C. Afterwards, cells were washed with Wash Buffer A (cat. #SMF-WA1-60, Biosearch technologies) for 5 minutes. Then the miR-200c-3p, Dll4 mRNA and Notch1 mRNA probes were mixed with the hybridization buffer (cat. # SMF-HB1-10, Biosearch technologies) according to the manufacturer’s instructions, covered in foil and placed in the cell incubator at 37°C for 5 hours, all subsequent steps were performed in the dark. Then, cells were incubated in Wash Buffer A for 30 minutes and placed in the incubator. Lastly, cells were incubated with Wash Buffer B (cat. # SMF-WB1-20, Biosearch technologies) for 5 minutes. Wells were replenished with fresh 1x PBS.

### 2.6 Imaging and data analysis

All images were acquired using a laser scanning confocal microscope (Leica TCS SP8; Leica Microsystems, Wetzlar, Germany). Wound healing migration data were analyzed in ImageJ. For all other experiments, the bright-field image was used as a template to create a mask to segment the images into single cells using Adobe Photoshop. The segmented images were then transferred to MATLAB for a complete analysis of morphological and fluorescence features at the single cell level.

### 2.7 Leader-follower cell selection and quantification

In this study, leader cells in the migrating monolayer are defined as cells at the migrating tip with apparent cell-cell contact with follower cells behind them. To be classified as a leader cell, we considered the distance from the initial wound boundary, the extent of the migration sprout, or tip, created by the leader cell, and the contact with follower cells. Follower cells were classified as those maintaining direct contact with the leader cell. To quantify number of leader cells per case (i.e., KO, WT, and SFN), the number of leader cells was counted per migration edge per case. The value was reported as leaders/mm, that is: (total # of leader cells / 1 mm wound edge).

### 2.8 Statistical analysis

Data obtained from MATLAB and ImageJ were analyzed using the statistical software GraphPad Prism 9. Experiments measuring mRNA/microRNA levels were performed at least 3 times in multiple experiments. All other assays were performed at least 4 times in multiple experiments. In single cell measurement experiments, at least 500 cells per case were analyzed. For first cell layer (i.e., the migrating edge) analyses, at least 100 cells per case were measured. All datasets were considered to follow a non-normal distribution. Therefore, non-parametric tests were utilized to compare across groups where possible. The tests used were: Kruskal-Wallis test with the Dunn’s multiple comparisons test, a Two-Way ANOVA test with a post-hoc Tukey test including multiple comparisons across rows and columns, and the ROUT method to identify outliers (Figures 3D and 7B). The following values were assigned to test for significance: ns p-value > 0.05, * p-value < 0.05, ** p-value < 0.01, *** p-value < 0.001, and **** p-value < 0.0001.

### 2.9 Multicell model of Nrf2-EMT-Notch signaling

We employed a continuous mass action model to describe the biochemical interactions between molecular players in the EMT, Nrf2 and Notch circuits. This approach was previously applied to the core regulatory circuits regulating EMT, Nrf2 and Notch signaling separately [17,29,33]. Within a cell, the temporal dynamics for the copy number of any given variable (say, ***X***) is described by the generic equation:

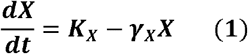

In eq. (1), the first term on the right-hand side (RHS) is a production rate with units of number of molecules produced per unit time. The effect of other microRNAs or transcription factors (TF) that induce or inhibit the production of ***X*** is modeled by additional functions that modulate the basal production rate. The second term on the RHS models molecule degradation. The full set of equations describing the EMT, Nrf2, Notch circuits and their mutual connections is presented in the supplementary sections A1-3.

The effect of transcriptional activation or inhibition exerted by a regulator (say, R) on another given species in the circuit is modeled with a shifted Hill function:

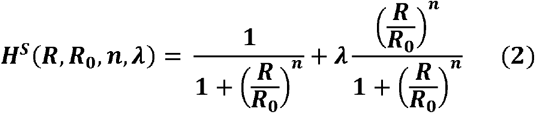

where ***R*** is the concentration or copy number of the regulator and ***R*_0_** is the half-maximal concentration parameter expressed in same units of ***R***. Additionally, the Hill coefficient ***n*** describes the steepness in transcriptional response with respect to the regulator concentration. Finally, the fold change ***λ*** describes the change in target level due to regulation by ***R*** (***λ*** < **1** implies that ***R*** is an inhibitor, while ***λ*** > **1** implies that ***R*** is an activator). If a species is regulated by multiple TFs, Hill functions are multiplied in the production rate of eq. (1).

Moreover, microRNAs can inhibit the production of other species in the circuit by binding the target mRNA and facilitating their degradation. This post-translational inhibition is modeled following the microRNA-TF chimera toggle switch model first introduced by Lu and collaborators [4]. First, a first set of functions ***P_l_***(***μ***, ***n***) quantifies the inhibition that a microRNA (***μ***) exerts on the target TF; here, ***n*** is the number of sites for microRNA binding to the mRNA. Furthermore, a second set of functions ***P_y_***(***μ***, ***n***) describes the corresponding decrease of microRNA due to the degradation of the microRNA/mRNA complex. The explicit form and derivation of these functions is discussed in the supplementary section A4.

In the multicellular model, cells are arranged on a two-dimensional hexagonal grid. The intracellular signaling dynamics of Nrf2, EMT and Notch is described within each cell by the coupled system of ODEs. Moreover, the biochemical networks of neighboring cells are coupled via ligand-receptor binding between Notch and its ligands, Dll4 and Jagged1. For any given cell (***i***) in the lattice, the numbers of external Notch receptors and ligands available for binding (***N_EXT_***^(***i***)^, ***D_EXT_***^(***i***)^, ***J_EXT_***^(***i***)^) are calculated as the sums over the cell’s nearest neighbors:

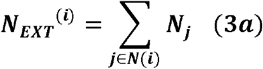

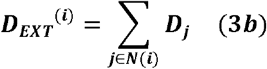

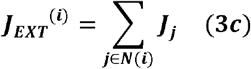

Moreover, to simulate the conditions of high stress close to the migrating front, we introduce a gradient of an EMT-inducing signal (***T***(***x***, ***y***, ***t***)) that is secreted at the left end of the lattice (the wound edge), diffuses along the x-coordinate and is removed at the opposite end of the lattice:

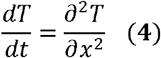

This diffusion dynamics gives rise to a profile where cells close to the invasive edge are highly exposed to EMT-inducing signaling while cells in the interior are weakly exposed. Simulation details and complete set of model’s parameters are provided in the supplementary section A5 and supplementary tables 2-5.

## 3. Results

### 3.1 Nrf2 modulates EMT during collective cell migration

We first evaluated the relationship between Nrf2 and EMT during the collective migration of cancer cells. Previous investigations in static cell monolayers using RT4 bladder cancer cells suggest that Nrf2 upregulation enhances the expression of both epithelial and mesenchymal markers (e.g., E-cadherin and ZEB1) while Nrf2 downregulation results in the attenuation of both markers [17]. To study the relationship between EMT and Nrf2 in migrating monolayers, we performed a scratch wound healing assay. Cells were then fixated and fluorescently labeled for Nrf2, E-cadherin, and ZEB1 after 24 hours (Figure 1A-C). The resulting images were then segmented into single cells for further analysis. For each gene, data were normalized to wild-type (WT) (Figure 1D-F). The complete cell array was analyzed and the mean intensity values were obtained for each cell to obtain an average intensity over the entire migrating monolayer (Figure 1D). Then, data were separated into cell layers (i.e., position relative to the migrating front) to study the spatial distribution near the leading edge (Figure 1E). The intensity distribution at the leading edge itself was further analyzed at the single cell level (Figure 1F).

**Figure 1.**
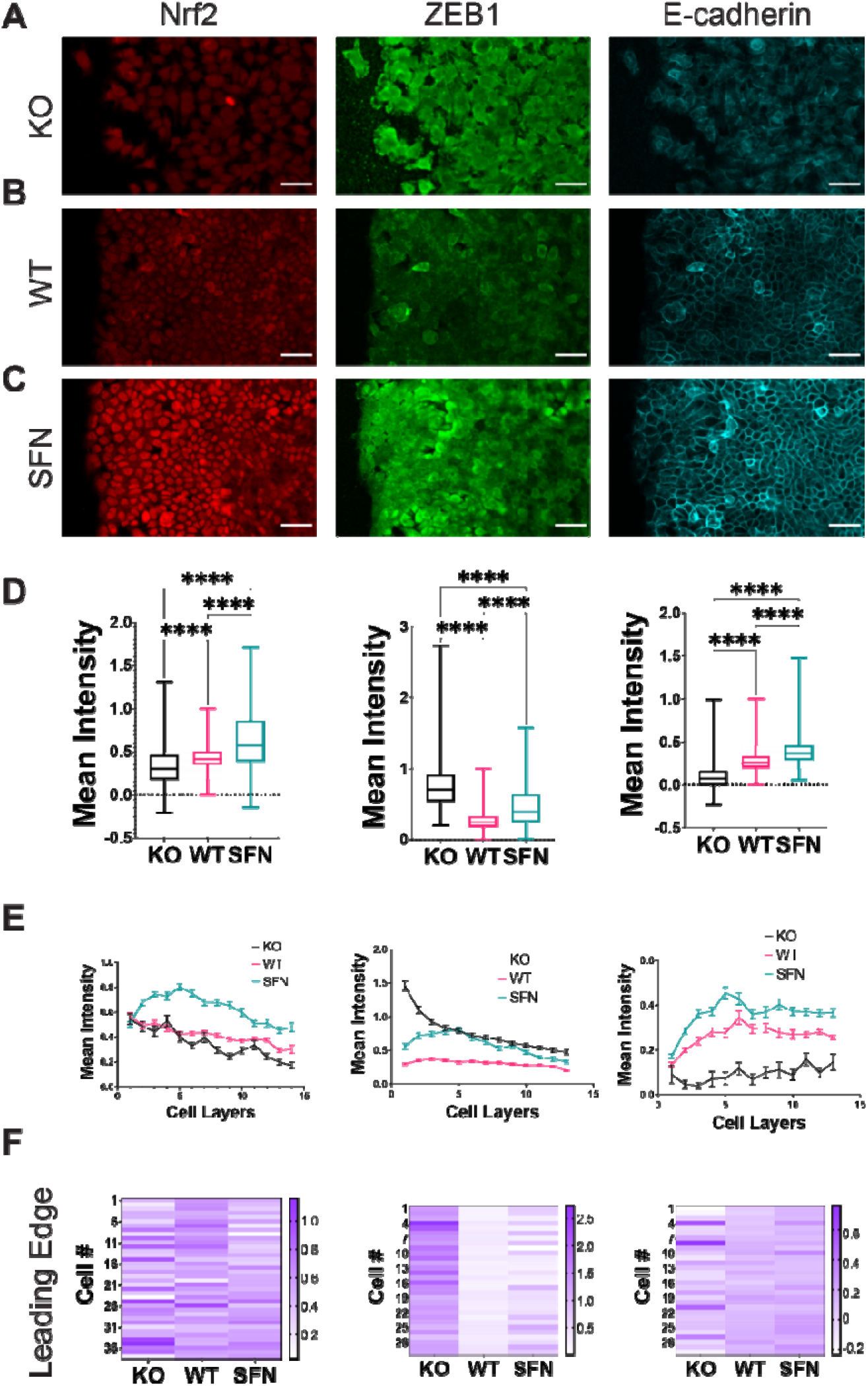
Nrf2 modulates EMT near the leading edge during collective migration. (A-C) Immunocytochemistry of RT4 bladder cancer cells measuring Nrf2 (left column), ZEB1 (center column), and E-cadherin (right column) protein levels in a wound healing assay for (A) CRISPR/Cas9 NFE2L2-KO Pool RT4 cells (KO), (B) wild-type RT4 cells (WT), and (C) sulforaphane treated (7.5 μM, 24h) RT4 cells (SFN), respectively. Scale bars, 50 μm. (D-F) Quantification of immunocytochemistry data. (D) Average intensity over the whole cell layer by measuring mean intensity of each cell in the whole monolayer, (E) intensity distribution in the migrating monolayer measured as number of cell layers (position from the leading edge), and (F) heatmap of representative cells near the wound leading edge for Nrf2, E-cadherin, and ZEB1, respectively. Each cell at the leading edge is indicated by “Cell #” where cell #1 refers to the first measured cell from top to bottom. The nonparametric Kruskal-Wallis test along with the Dunn’s multiple comparisons test were used to compare across groups. For each experiment n > 500 cells per condition. **** p-value < 0.0001. Images are representatives from 6 experiments.

Nrf2 upregulation via sulforaphane (SFN) treatment and Nrf2 downregulation in CRISPR-Cas9 RT4-Pool-knockout (KO) cells resulted in significant enhancement and reduction of Nrf2 across the migrating monolayer, respectively (Figure 1D, left column). Moreover, the Nrf2 distribution showed a significant dependence on the distance from the migrating front. Sulforaphane treatment enhanced the overall level of Nrf2, as expected, but also shifted the maximum level of Nrf2 toward the interior region (layers 3-7) of the migrating cell monolayer (Figure 1E, left column). In contrast, Nrf2 KO resulted in a significant decline in the Nrf2 expression, especially in the interior region of the migrating monolayer. We observed no significant difference across groups at the leading edge (Figure 1F, left column).

We further examined the EMT markers in the migrating monolayer. In the WT case, a reduction of E-cadherin was observed near the leading edge, suggesting that cells at the migrating front may undergo EMT, reminiscent of earlier observations [14]. For the KO group, we observed an overall reduction of E-cadherin and an enhancement of ZEB1, thus suggesting that cells displayed a more mesenchymal phenotype as compared to WT. Both WT (control) and Nrf2 KO exhibited EMT activation that gradually fades as a function of distance from the leading edge. In contrast, the sulforaphane group showed high levels of both E-cadherin and ZEB1 compared to the WT control and thus suggesting a hybrid E/M phenotype (Figure 1D, center and right columns). Furthermore, when examining the spatial distribution, E-cadherin was lowest near the wound edge across all groups, and ZEB1 was highest near the wound edge for the KO group. Interestingly, both E-cadherin and ZEB1 were maximized at rows 3-7 for the sulforaphane group (Figure 1E-F, center and right columns). The formation of hybrid E/M cells was further analyzed by estimating the intensity product of E-cadherin and ZEB1 (Supplementary Figure S2). The intensity product, which signifies cells with both mesenchymal and epithelial signatures, was maximized in the interior region (~ row 5) of the migrating monolayer for WT and sulforaphane. This value was enhanced with sulforaphane treatment, and the peak shifted toward the leading edge. These results suggested that Nrf2 prevents a complete EMT and instead stabilizes a hybrid E/M cell phenotype near but not directly at the leading edge during collective cancer migration.

### 3.2 In silico modeling predicts Nrf2-dependent increase of the hybrid E/M cell population near the leading edge

To gain further insight into the role of Nrf2 in regulating EMT, we turned to *in silico* modeling of the underlying regulatory dynamics. We have previously developed circuit models governing EMT-Nrf2 intracellular crosstalk as well as EMT-Notch multicellular signaling dynamics [17,29,33]. These models predicted that Nrf2 expression stabilizes a hybrid E/M phenotype. Here, we integrated these models into a more comprehensive framework to investigate how cell-cell and cell-environmental interactions in a wound-healing scenario modulate the connection between Nrf2 signaling and EMT. In the computational model, the biochemical dynamics within each cell is described by interconnected feedbacks between the Nrf2, EMT, and Notch signaling modules. Moreover, binding between Notch1 and its ligands (Dll4 and Jagged1) enables communication between the biochemical circuits of neighboring cells (Figure 2A, right). Simulated cells were exposed to an EMT-inducing signal that diffused throughout the cell layer, thus allowing our model to mimic the spatially-dependent cellular response to the presence of the wound edge. Thus, cells toward the invasive edge (the leftmost side) of the lattice are highly exposed to an EMT-inducing signal, while cells in the interior (the rightmost side) of the lattice are only weakly exposed (Figure 2A, left). Varying the EMT inducer level modules the level of EMT. The invasive edge can be mostly composed by mesenchymal cells at high EMT induction or by mixed E/M and epithelial cells at low EMT induction (Supplementary Figure S3).

**Figure 2.**
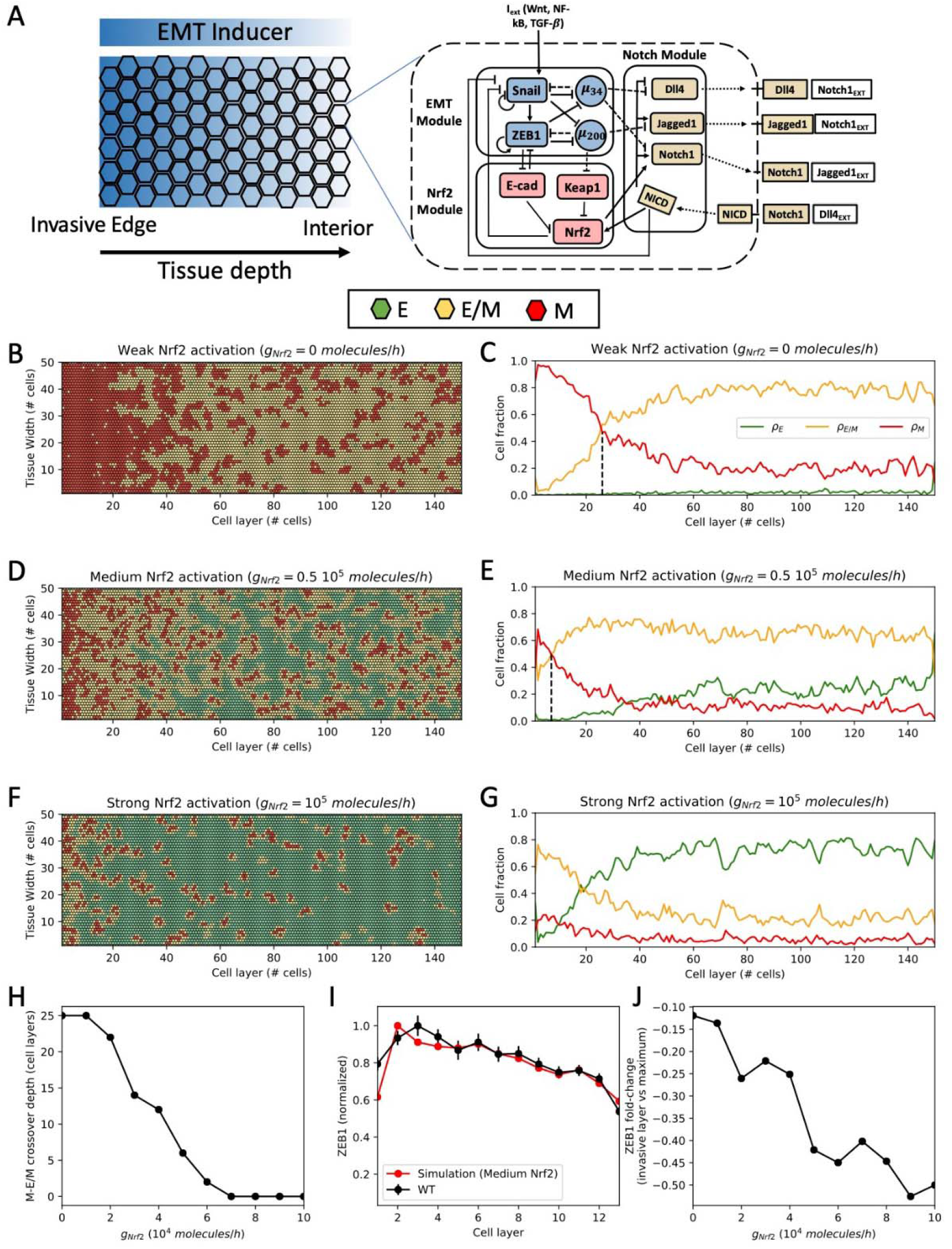
Spatial patterning of cells in the multicell model of wound healing. (A) Left: In the multicell model, cells are arranged on a hexagonal lattice. Cells at the leftmost region (invasive edge) are highly exposed to an external EMT-inducing signal, while cells in the interior are weakly exposed. Right: The signaling dynamics within each cell is described by the coupled biochemical network of Nrf2, EMT and Notch. Binding between Notch ligands and receptors of neighboring cells give rise to cell-cell communication. (B) Snapshot of the multicell pattern after 120 hours of simulation starting from a randomized initial condition for a case of weak Nrf2 activation (molecules/hour). Green, yellow and red hexagons depict epithelial (E), hybrid E/M and mesenchymal (M) cells, respectively. (C) Fraction of E, E/M, and M cells as a function of distance from the leading edge. (D-E) Same as (B-C) for intermediate Nrf2 activation (molecules/hour). (F-G) Same as (B-C) for strong Nrf2 activation (molecules/hour). (H) Crossover point where the fraction of hybrid E/M cells becomes larger than the fraction of M cells as a function of Nrf2 production rate. (I) Comparison of ZEB1 levels between simulation (red line) and WT experiment (black line) in the first 13 cell layers. (J) Fold-change in ZEB1 levels between the first and second cell layers as a function of Nrf2 production rate.

By treating the production rate of Nrf2 as a controllable parameter, we investigated the cell layer’s response to varying levels of Nrf2 induction. In the model, cells are categorized as epithelial, hybrid E/M, or mesenchymal based on their decreasing levels of the epithelial microRNA miR-200 [4]. Starting from randomized initial conditions, cell populations evolve in time depending on Nrf2 induction and distance from the wound edge.

At the basal, or medium, level of Nrf2, the first 5-10 cell layers were mostly composed by mesenchymal cells, while the more interior region was mainly composed by hybrid E/M and epithelial cells (Figure 2D and Supplementary Movie 1). A weaker Nrf2 induction increased the mesenchymal cell population at the migration front while pushing the hybrid E/M cell population to a more interior region of the monolayer (Figure 2B-C and Supplementary Movie 2). Furthermore, the epithelial phenotype was almost completely suppressed. Therefore, a change from medium to weak Nrf2 induction in the computational model recapitulates many of the experimental findings seen when comparing the WT with Nrf2 KO cases. In contrast, for a strong Nrf2 induction, the hybrid E/M cell population became dominant even at the leading edge, in good agreement with the high expression of both ZEB1 and E-cadherin in the experimental sulforaphane case (Figure 2F-G and Supplementary Figure S2).

The phenotype distribution can be quantified by a ‘crossover point’, where the hybrid E/M cell fraction becomes larger than the mesenchymal cell fraction (dashed lines in Fig. 2C and E). This transition point depends on several model’s parameters, including the concerted effect of cell-autonomous EMT-induction driven by the signaling gradient and cell-cell communication EMT-induction driven by Notch signaling (Supplementary Figure S4). Nrf2 induction, however, moved the crossover point toward the invasive edge independently of the other model’s parameters (Fig. 2H), thus supporting the role for Nrf2 in preventing complete EMT and stabilizing the hybrid E/M phenotype near the migrating front of cells.

The model predicts a drop in the mesenchymal cell fraction at the invasive edge, which is instead maximized in the cell layers immediately behind (see for instance Fig. 2C, 2E). Invading cells at the free end received Notch-mediated EMT induction from a smaller number of neighbors, thus resulting in more hybrid E/M cells and less mesenchymal cells. Remarkably, a drop in ZEB1 levels at the leading edge was also observed in both WT and SFN experiments (P<0.001), and can be at least semi-quantitatively compared to the simulation results (Fig. 2I). Conversely, the KO experiment did not exhibit a ZEB1 drop. It can be speculated that Notch signaling plays a lesser role due to the loosen adhesive bonds between the highly mesenchymal cells observed in the KO case, a feature that is not captured by the current model. Moreover, the ZEB1 drop between first and second cell is predicted to increase as a function of Nrf2 (Fig. 2J). This trend is qualitatively observed in the experimental model as well, where the SFN case presents a larger drop compared to the WT. Overall, the computational model suggests a role for Nrf2 in modulating the spatial composition of the cell layer by preventing a complete EMT and localizing a population of hybrid E/M cells at the migrating edge, in good agreement with high co-expression of epithelial and mesenchymal markers observed under sulforaphane treatment. The coupled dynamics between Nrf2, EMT and Notch signaling drives Nrf2 to act as a brake on EMT, thus increasing the population of hybrid E/M cells near the wound edge.

### 3.3 Nrf2 modulates Notch signaling near the leading edge

The spatial patterning determined via our computational model depends directly on cell-cell coupling via the Notch pathway. From an experimental perspective, Notch signaling has been shown to be a critical component of EMT circuitry [34,35]. Also, Notch signaling has been shown to regulate collective cell migration [22–26]. We therefore measured the distributions of Notch components (i.e., Notch1, Dll4, and Jagged1) in the WT and under Nrf2 perturbations (Figure 3A-C). In agreement with previous studies [22,25,26], Notch signaling was upregulated near the leading edge. Spatial gradients of Notch1, Dll4, and Jagged1 near the leading edge were observed, and the expression levels were dependent on Nrf2. In particular, Jagged1 was negatively correlated with Nrf2 levels, being consistently highest for the KO case and lowest in the sulforaphane case, in the entire monolayer (Figure 3D-F, left column). Notch1 and Dll4 showed mutually exclusive behavior (as expected), but did not respond proportionally to Nrf2 induction. Specifically, Notch1 was lowest for the WT case and higher for both KO and sulforaphane, whereas Dll4 was highest for the WT case and lowest for KO and sulforaphane cases (Figure 3D, center and right column). This behavior was especially apparent when inspecting cells near the leading edge (e.g., the first 5 rows), where we noticed a great degree of separation between the WT behavior and that of the other two cases (Figure 3E, center and right column). At the leading edge, Notch1 showed lowest intensity values in the WT case whereas Dll4 showed the highest intensity values for the WT case (Figure 3F, center and right column).

**Figure 3.**
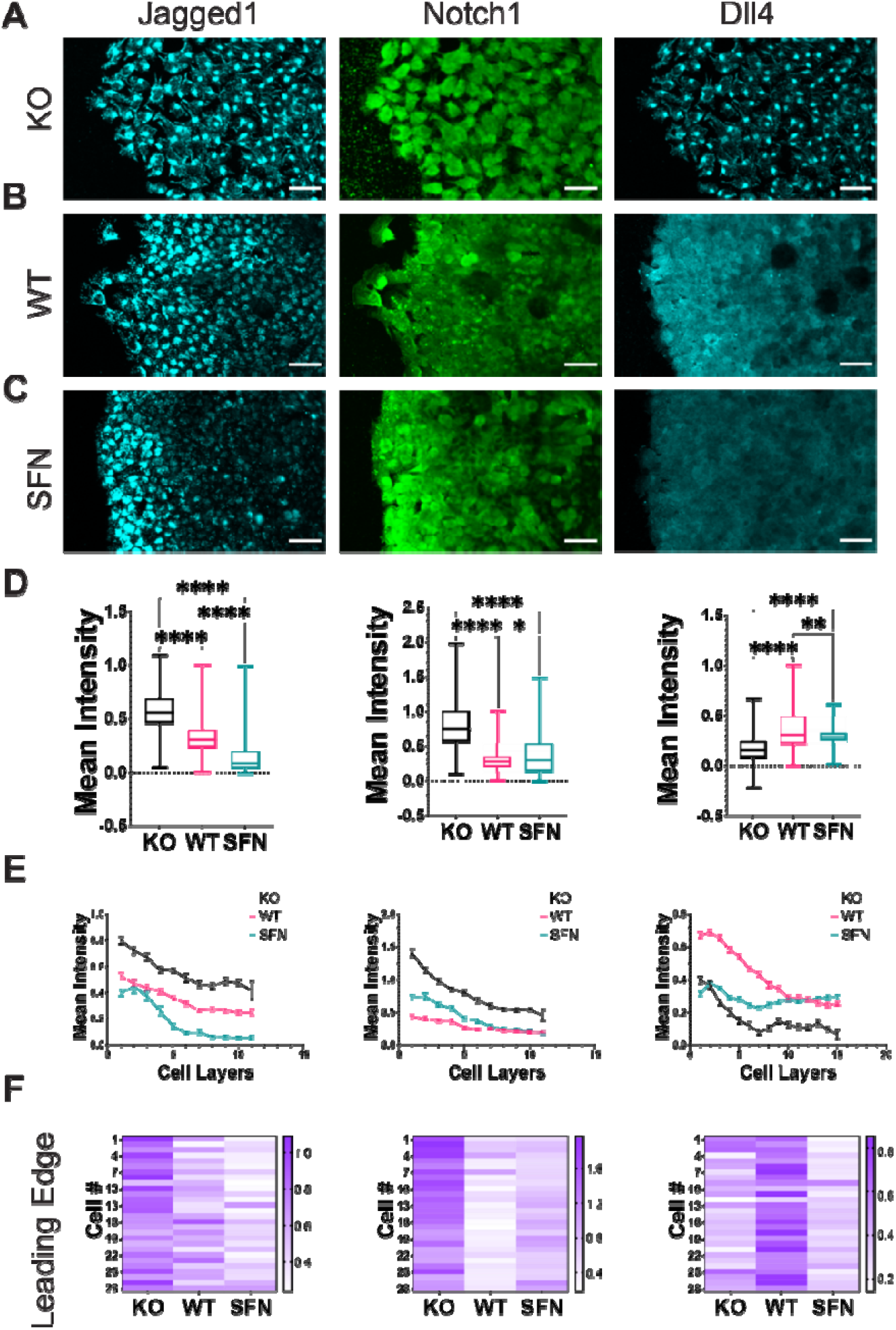
Nrf2 regulates the Notch signaling pathway near the wound edge. (A-C) Immunocytochemistry of RT4 bladder cancer cells measuring Jagged1 (A-C left column), Notch1 (A-C center column), and Dll4 (A-C right column) protein levels in a wound healing assay for (A) CRISPR/Cas9 NFE2L2-KO Pool RT4 cells (KO), (B) wild-type RT4 cells (WT), and (C) sulforaphane treated (7.5 μM, 24h) RT4 cells (SFN), respectively. Scale bars, 50 μm. (D-F) Quantification of immunocytochemistry data. (D) Average intensity over the whole cell layer, (E) tracing of relative fluorescence intensity per tissue depth measured as number of cell layers, and (F) heatmap of representative cells near the wound leading edge for Jagged1, Notch1, and Dll4, respectively. Each cell at the leading edge is indicated by “Cell #” where cell #1 refers to the first measured cell from top to bottom. The nonparametric Kruskal-Wallis test along with the Dunn’s multiple comparisons test were used to compare across groups (view Materials and Methods section). For each experiment n > 500 cells per condition. ns: not significance, * p-value < 0.05,** p-value < 0.01, and **** p-value < 0.0001.

Dll4 mRNA has been reported to be upregulated in leader cells during collective migration [22,25,26]. Particularly, mRNA levels of Dll4 are upregulated in leader cells and exhibit a higher contrast between leader and follower cells when compared to Dll4 protein [25]. We therefore directly evaluated the influence of Nrf2 activation on the expression of Notch1 mRNA and Dll4 mRNA. We also measured the miR-200c-3p, which is a key component of the regulatory circuit driving hybrid E/M and can attenuate Jagged1 [29]. Specifically, we used a double-stranded single cell biosensor as well as the FISH assay to measure the expression levels of miR-200c-3p, Notch1, and Dll4 near the leading edge (Figure 4). Biosensors were added prior to the wound healing assay to ensure uniform probe internalization [32]. Images of live migrating cells were acquired 24 hours post wound scratch to characterize the gene expression. The left panel shows fluorescence images for WT and sulforaphane cases measuring miR-200c, Notch1, and Dll4 with the single cell biosensor (Figure 4A-C, left panel) and in the FISH assay (Figure 4D-F, left panel). The right panels indicate the intensity distribution as a function of distance from the migrating front and a representative distribution at the leading edge (Figure 4A-F, right panel).

**Figure 4.**
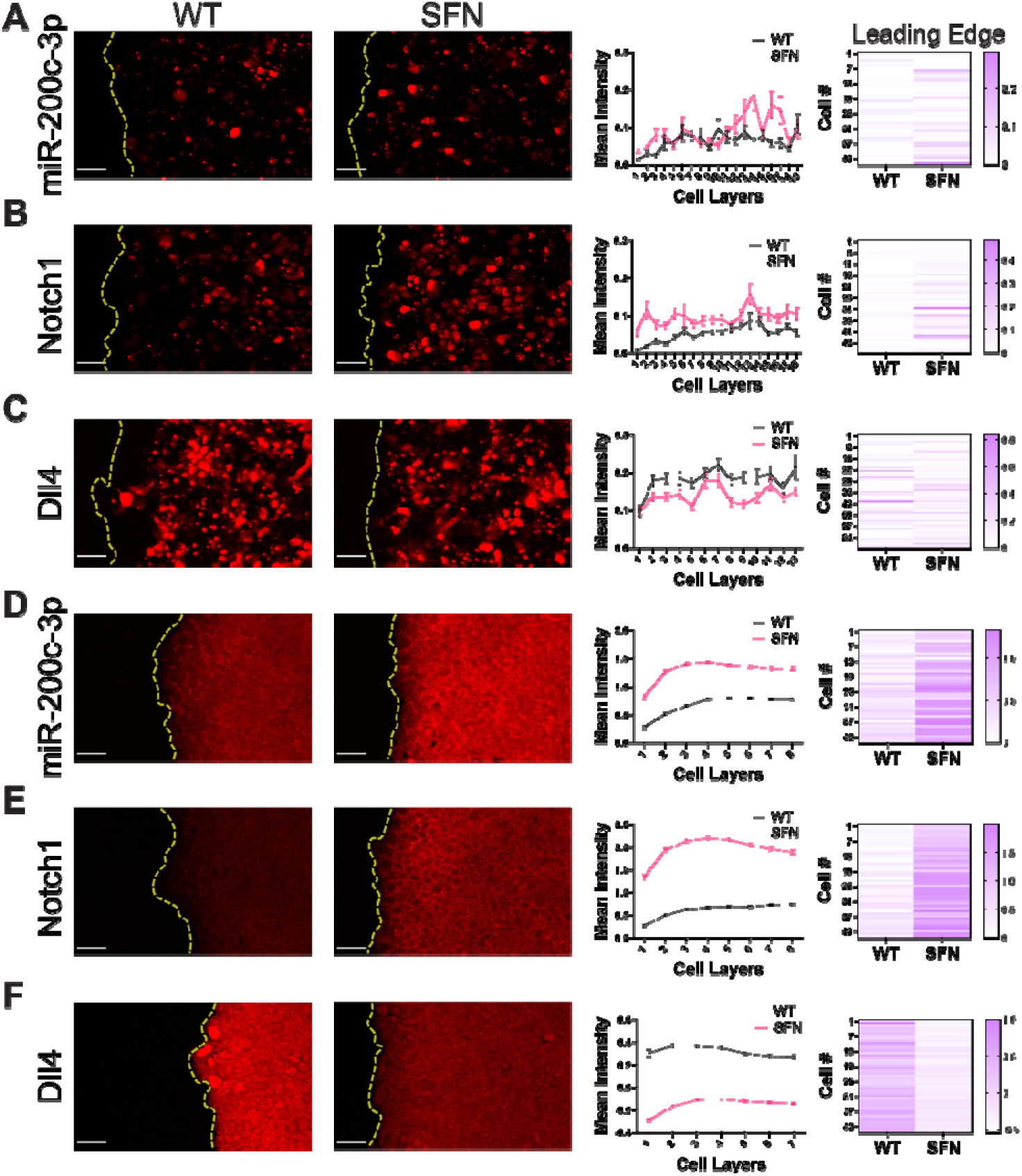
Nrf2 upregulation modulates microRNA miR-200c-3p and Notch1 and Dll4 mRNA near the wound edge. (A-C) Live single cell gene expression measurements with the dsLNA probes in RT4 cells. From left to right: fluorescence images of WT and SFN (7.5 μM, 24h) cases, tracing of relative fluorescence intensity per tissue depth measured as number of cell layers, and heatmap of representative cells near the wound leading edge measuring (A) microRNA miR-200c-3p, (B) Notch1 mRNA, and (C) Dll4 mRNA levels in a scratch wound healing assay. (D-F) FISH assay in fixed RT4 cells. From left to right: fluorescence images of WT and SFN cases, tracing of relative fluorescence intensity per tissue depth measured as number of cell layers, and heatmap of representative cells near the wound leading edge measuring (D) microRNA miR-200c-3p, (E) Notch1 mRNA, and (F) Dll4 mRNA levels in a scratch wound healing assay. Scale bars, 50 μm. Each cell at the leading edge is indicated by “Cell #” where cell #1 refers to the first measured cell from top to bottom. The 2-way ANOVA with the post-hoc Tukey test to compare across groups was performed as well as the nonparametric Mann-Whitney test. For each experiment n > 500 cells per condition.

A gradient of miR-200c-3p was observed near the migrating front. The level of miR-200c-3p was lowest at the leading edge and increased toward the interior region, consistent with the spatial gradient observed in E-cadherin immunostaining. Furthermore, Nrf2 activation by sulforaphane treatment enhanced the level of miR-200c-3p corresponding to an increase in epithelial and hybrid E/M cells. The gradient of miR-200c-3p and the influence of Nrf2 activation were in good agreement with the predictions of the computational model. Furthermore, Nrf2 activation suppressed the average level of Dll4 and enhanced Notch1, similar to the immunocytochemistry analysis. Notably, non-uniform distributions of Dll4 were observed at the leading edge, especially for the WT case. In particular, a small number of cells at the leading edge displayed a high level of Dll4. As discussed below, these cells can be identified as leader cells during collective cell migration.

### 3.4 Computational modeling predicts a NRF2-dependent transition in Notch signaling mode and EMT, at the leading edge

Next, we returned to the mathematical model to investigate whether the detailed response of Notch1, Dll4 and Jagged1 to Nrf2 modulation could be understood in terms of the interconnected feedbacks between the Notch1 and Nrf2 pathways. Since we were especially interested in Nrf2’s role in mediating collective migration and leader cell formation, we focused our analysis on the leading edge of the multicell model and created simulated data that could be directly compared to the front of migrating monolayer (Figure 5A). As Nrf2 induction increased, the molecular composition of the leading edge changed substantially. Nrf2 induction increased the number of cells with high levels of miR-200 and Notch1 (Figure 5B). The increase of Notch1 can be understood by the mutual positive feedback between NICD, Nrf2, and the Notch receptor (see circuit in Figure 2A). Notably, low-Notch cells were still observed for high Nrf2 induction levels due to lateral inhibition promoted by Notch1-Dll4 signaling. Nrf2 induction also decreased the frequency of cells with high Dll4 and high Jagged1 (Figure 5B). In the case of low Nrf2 induction, most cells either expressed high Dll4 or high Jagged1. Interestingly, a small fraction of cells co-expressed both ligands, reflective of a hybrid Sender/Receiver phenotype enabled by a balance of lateral inhibition and lateral induction (Figure 5C) [36]. This effect was progressively removed by a stronger Nrf2 induction, as cells expressing either one or both ligands become rarer (Supplementary Figure S5A-C).

**Figure 5.**
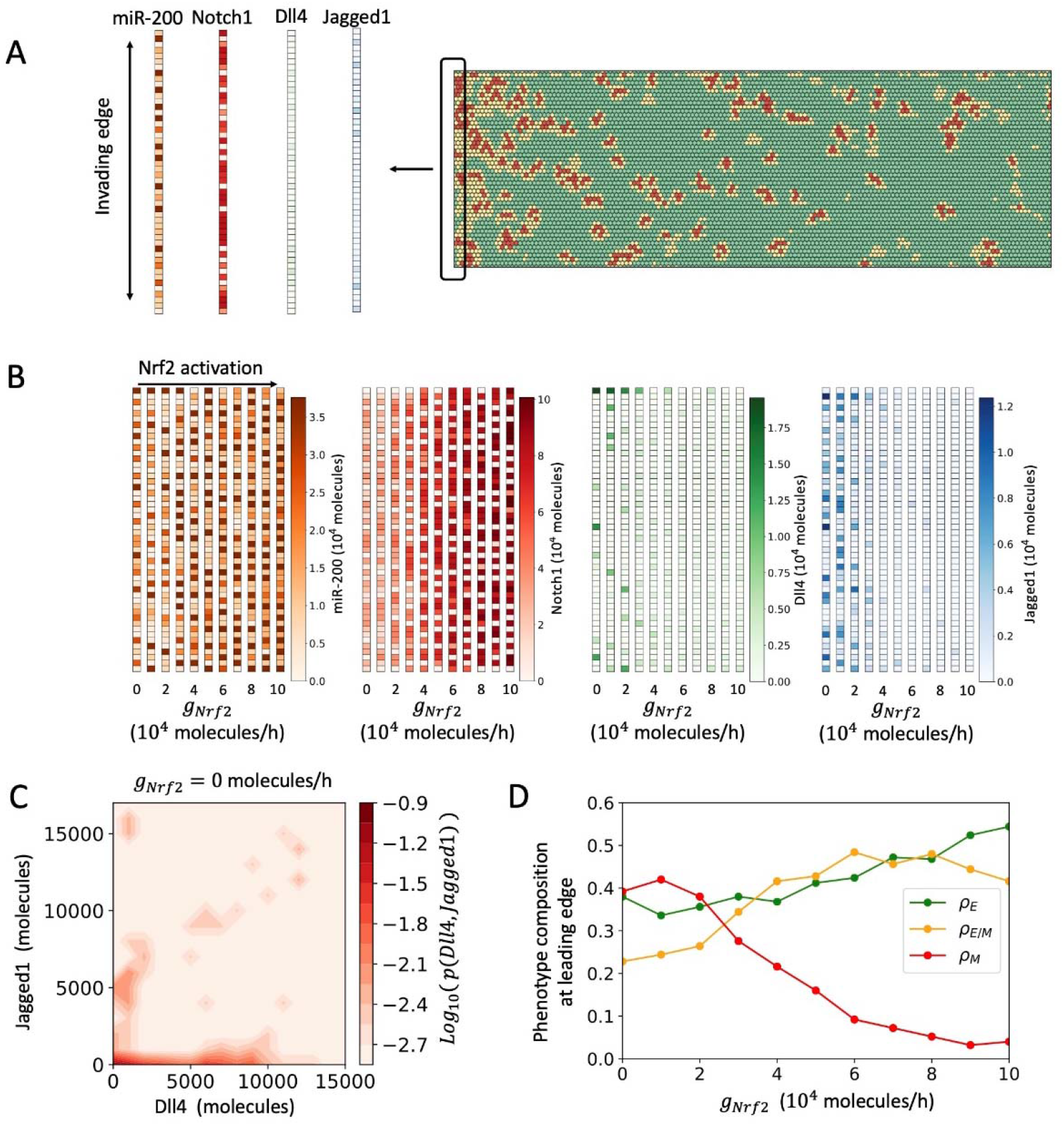
Analysis of the invading edge by the multicell model. (A) To conduct invading layer analysis, the expression of miR-200, Notch, Dll4 and Jagged1 is analyzed in the leftmost layer of cells (i.e., the cell layer more exposed to EMT-inducing signal). (B) Heatmap of expression levels for miR-200, Notch1, Dll4 and Jagged1 in the invading layer as a function of Nrf2 activation. (C) Log-normalized probability to observe cells with varying levels of Dll4 and Jag4 in the invasive edge under low Nrf2 induction. (D) Fraction of Epithelial, hybrid E/M, and Mesenchymal cells in the leading edge as a function of Nrf2 activation. For panels C-D, results are averaged over 5 simulations starting from randomized initial conditions.

In terms of EMT phenotype composition, the invading edge was predominantly composed of mesenchymal cells when Nrf2 induction is low. Conversely, at higher Nrf2 levels, the invasive edge was a mixture of hybrid E/M and epithelial cells (Figure 5D). Varying the relative strength of Notch1-Dll4 and Notch1-Jag1 signaling modulates the composition of the leading edge. However, Nrf2 induction restricted the fraction of mesenchymal cells while increasing the fraction of hybrid E/M cells (Supplementary Figure S5D-G). More generally, Nrf2 induction correlated with an average increase in miR-200 and Notch1 expression at the invasive edge, as well as decrease of Dll4 and Jagged1 expression, similar to the trend observed in the experiments from WT to SFN (Supplementary Figure S5H-I). The trends of Notch1, Dll4 and Jagged1 as a function of Nrf2 induction were confirmed also when inspecting the expression throughout the whole lattice model (Supplementary Figure S6). Noticeably the opposite, experimentally-observed trend of Notch1 and Dll4 from KO to WT cannot be reproduced, potentially suggesting that other factors besides Notch-Nrf2 interactions might modulate this response. Overall, the model predicts that Nrf2 induces a transition from a mostly mesenchymal invading edge with strong Dll4 and Jagged1 signaling to a mostly hybrid E/M and epithelial invasive edge with high Notch1 expression, in good agreement with the trend observed experimentally when increasing Nrf2 activation from WT to sulforaphane.

### 3.5 Leader cell formation is optimal for the WT case and Dll4 is highest near the wound edge

The modulation of the Notch ligands Dll4 and Jagged1, which are associated with leader cells [19], suggest Nrf2 may modify the formation of leader cells during collective cancer migration. We thus investigated leader cells at the wound edge (Figure 6). We defined leader cells based on their spatial location at the protruding tips and their interactions with follower cells. Bright-field images at the leading edge revealed distinct morphologies of leader cells for KO, WT, and sulforaphane cases (Figure 6A). When treated with sulforaphane, leader cells showed a less mesenchymal phenotype with smaller cell size compared to Nrf2 KO. Leader cells in the WT case showed aggressive morphologies, including enlarged cell size and active lamellipodial structures. Moreover, leaders in the WT case appeared to entrain a larger number of follower cells when compared to KO and sulforaphane cases (Figure 6A). The WT case also exhibited the highest density of leader cells when compared to the other cases (Figure 6B).

**Figure 6.**
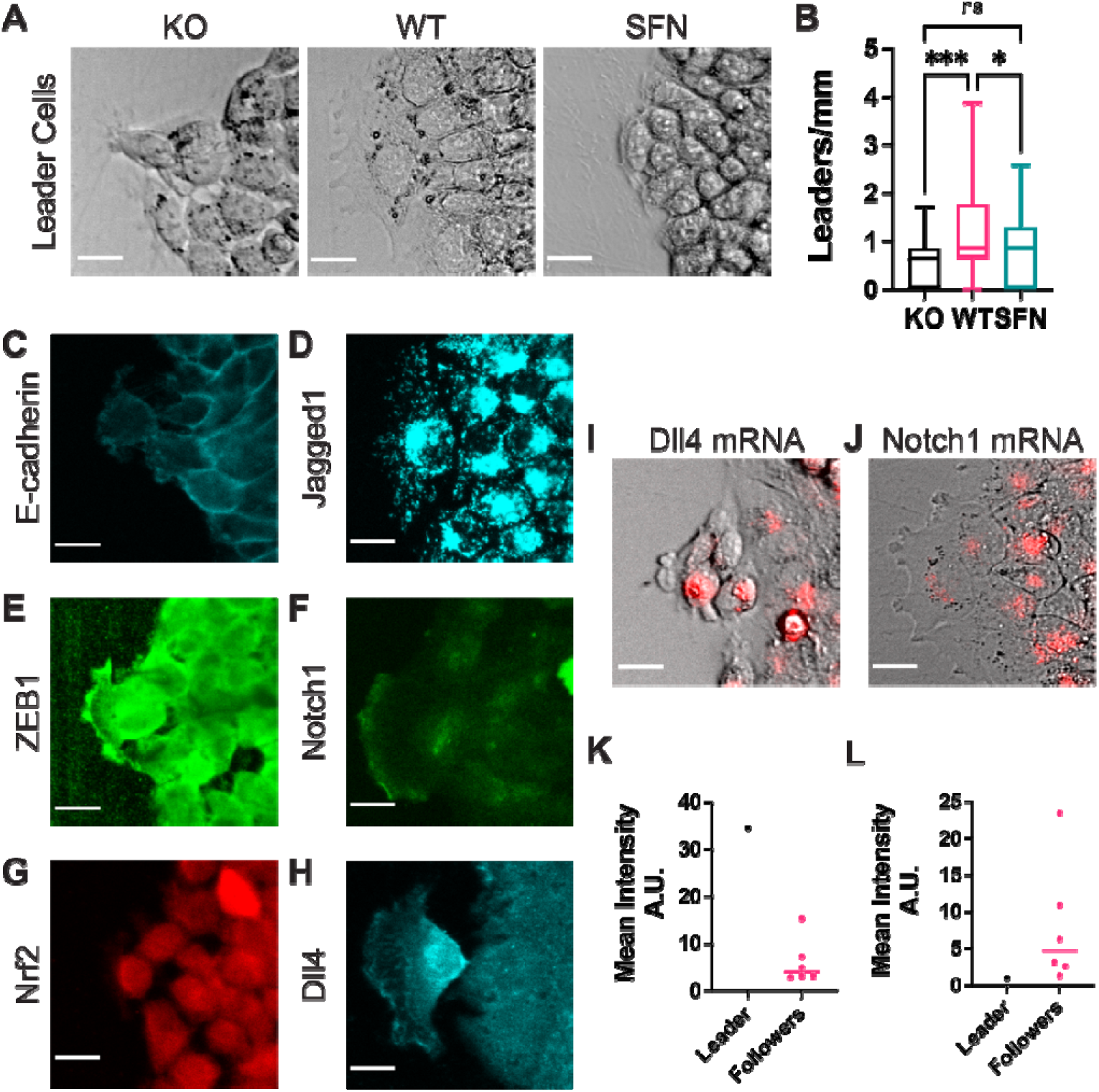
Leader cell formation near the leading edge. (A) Representative bright-field images of leader cells after a 24h wound healing assay for CRISPR/Cas9 NFE2L2-KO Pool RT4 cells (KO), wild-type RT4 cells (WT) and sulforaphane treated (7.5 μM, 24h) RT4 cells (SFN), respectively. (B) Bar chart showing leader cells per millimeter for CRISPR/Cas9 NFE2L2-KO Pool RT4 cells (KO), wild-type RT4 cells (WT) and sulforaphane treated RT4 cells (SFN), respectively. ns: not significance, * p-value < 0.05, and *** p-value < 0.001. (C-H) Representative immunocytochemistry images of WT leader cells characterizing gene expression for (C) E-cadherin, (D) Jagged1, (E) ZEB1, (F) Notch1, (G) Nrf2, (H) Dll4. (I-J) Representative composite images using the dsLNA biosensors to measure mRNA levels of (I) Dll4 and (J) Notch1, respectively. (K-L) Mean intensity of leader vs follower cells for levels of (K) Dll4 mRNA and (L) Notch1 mRNA. Scale bars, 20 μm. The nonparametric Kruskal-Wallis test along with the Dunn’s multiple comparisons test were used to compare across groups and the ROUT method was used to identify outliers in (B). For each condition n > 40 scratch wound experiments.

We further analyzed molecular markers of leader cells in the WT case at the protein and mRNA levels (Figure 6C-H). Leader cells generally showed a low level of E-cadherin and a high level of ZEB1. This is expected as most cells at the leading edge exhibit a mesenchymal phenotype. Leader cells also expressed a low level of Notch1 while upregulating both Jagged1 and Dll4. This observation is particularly interesting as Dll4 and Jagged1 are often assumed as mutually exclusive Notch signaling states [37,38], and agrees with the model’s prediction that a small population of high Dll4 and high Jagged1 cells exists at the leading edge due to biochemical feedbacks between the Notch, EMT and Nrf2 signaling modules. Furthermore, Jagged1 was relatively uniform in all cells at the leading edge while Dll4 was selectively upregulated in leader cells (Figure 6H). The selective upregulation of Dll4 in leader cells was particularly profound at the mRNA level (Figure 6I,K), which is consistent with previous leader cell investigation, where Dll4 was distinctively upregulated in leader cells at the mRNA level [25]. In turn, Notch1 was dramatically downregulated in leader cells while being upregulated in follower cells (Figure 6J,L). This is in agreement with model predictions indicating the mutually exclusive states for Dll4 and Notch1.

### 3.6 Nrf2 and collective cell migration

Lastly, we analyzed how Nrf2 affects the overall collective migration of cancer cells. The migration of the scratch wound was measured at 0 and 48 hours for all cases (Figure 7A). We observed a decrease in migration speed in both the KO and sulforaphane cases (Figure 7B). Specifically, the WT case was significantly faster than both the KO and sulforaphane cases (p < 0.0001, n > 40 cases). This trend correlated with Dll4 expression and the formation of leader cells. Furthermore, “protruding tips” were formed at the leading edge (Figure 7C). The protruding tips often consisted 10-20 cells extended beyond the wound boundary, resulting in an irregular wound edge. These protruding tips were most profound in the WT case (see also Figure 6A). The KO and sulforaphane cases, in contrast, displayed relatively uniform wound boundary and had few or smaller protruding tips (Figure 7D).

**Figure 7.**
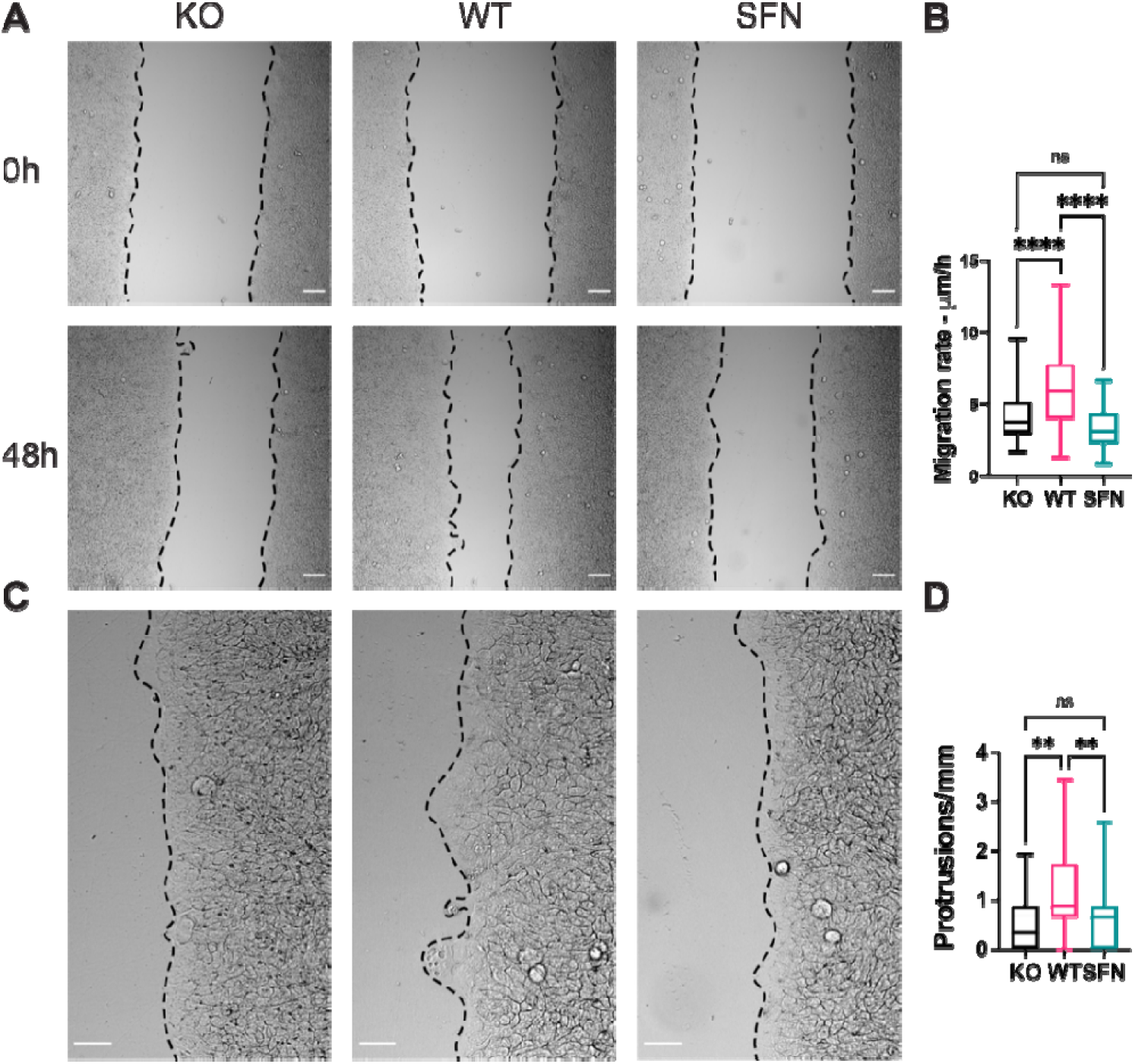
Nrf2 modulations impair collective migration in a 2D wound healing model. (A) Bright-field images of CRISPR/Cas9 NFE2L2-KO Pool RT4 cells (KO), wild-type RT4 cells (WT) and sulforaphane treated (7.5 μM, 24h) RT4 cells (SFN) for 0 h and 48 h migration time points. Scale bars, 100 μm. (B) Boxplot of migration rate for KO, WT, and SFN, respectively. (C) Representative images illustrating the formation of migration tips. Scale bars, 100 μm. (D) Boxplot of protrusion tips per millimeter for KO, WT, and SFN, respectively. The nonparametric Kruskal-Wallis test along with the Dunn’s multiple comparisons test were used to compare across groups and the ROUT method was used to identify outliers in (D). For each condition n > 40 scratch wound experiments. ns: not significance, ** p-value < 0.01, and **** p-value < 0.0001.

## 4. Discussion

This study investigated the role of Nrf2 in modulating the hybrid E/M state(s) of collectively migrating bladder cancer cells. While Nrf2 has been widely studied in the context of antioxidant response and chemoresistance [39,40], its function in cancer progression and invasion remains poorly understood [15,16]. Our experimental-computational analysis revealed that the Nrf2-EMT-Notch network coordinates cancer cells near the leading edge during collective migration. In particular, Nrf2 acted as a PSF in suppressing a full EMT and promoting a hybrid E/M phenotype, in a spatially coordinated manner. In the unperturbed condition (i.e., WT), the cells near the migrating front displayed a gradient of epithelial to mesenchymal behaviors. The cells near the leading edge were relatively mesenchymal while the cells in the monolayer displayed an epithelial phenotype. It should be noted that even cells at the leading edge maintained cell-cell contact with neighboring cells and expressed a detectable level of E-cadherin, supporting a partial EMT identification instead of a full EMT. In both experimental and computational models, Nrf2 stabilized the hybrid E/M cells, which were positioned at an interior region behind the leading edge. Nrf2 activation was required to maintain the hybrid E/M state of these cells. As indicated in Nrf2 KO, the cells shifted toward a more mesenchymal state, and the level of E-cadherin expression was significantly attenuated in the migrating monolayer. In contrast, Nrf2 upregulation enhanced the hybrid E/M phenotype and increased both epithelial and mesenchymal markers, especially in the interior region (several rows behind the leading edge) in the migrating monolayer.

Our data implicate a potential function of the Nrf2-EMT-Notch network in spatially coordinating collective cancer migration. In particular, cancer cells several rows behind the leading edge exhibited upregulated Notch1, Nrf2, and miR-200c while cells at the leading edge expressed ZEB1, Dll4, and Jagged1. The spatial coordination is contributed by the elevated Nrf2-Notch-NICD activity in the interior region and enhanced ZEB1, which reduced miR-200c [18] and consequently increased Jagged1, at the leading edge. Importantly, the Nrf2-EMT-Notch network promoted the upregulation of Dll4 and Jagged1 at the leading edge, which appeared to correlate with the formation of leader cells and protrusion tips. Dll4 is associated with the formation of leader cells during collective cancer invasion [22,25,26]. Jagged1 is also shown to promote MYO10 driven filopodial persistence for fibronectin micropatterning of leader cells [28]. Our computational analysis shows that the coordination between Dll4 and Jagged1 is highly sensitive to Nrf2 activation. Our predictions indicate that cells expressing both Dll4 and Jagged1 should exist at the leading edge (Figure 6C). Therefore, our results suggest Nrf2 may play a role in the coordination of Dll4 and Jagged1 near the leading edge to regulate different aspects of leader cells. Similar hypotheses have been drawn in contexts of inner ear development and sprouting angiogenesis, where a weak Jagged1 signaling was proposed to further amplify Notch1-Dll4 lateral inhibition by further sequestering Notch1 ligands in high Notch1 receiver cells [38,41]. Moreover, Jagged1 are also implicated in inducing partial EMT and cancer stem cell traits and propagating these aggressive traits to neighboring cells via Notch1-Jagged1 signaling [38,41,42]. The precise dynamics acting between Dll4 and Jagged1 and its functional implications in cancer invasion should be further investigated.

Our data indicate that the overall migration speed of collective cancer migration is sensitive to changes in Nrf2 activity. Previous studies report both positive and negative effects of Nrf2 on the collective invasion of various cancer cell types [43–47]. In general, cancer cell migration can be influenced by multiple factors, such as leader cell formation, cell motility, and proliferation, which can be modulated by Notch and EMT signaling both directly and indirectly. Cancer cells, at least in our model of urothelial bladder cancer, have their maximum migration speed in the unperturbed condition. The migration speed correlated with Dll4 expression and the formation of protruding tips and leader cells. Furthermore, EMT is associated with the motility and proliferation of cancer cells [2,3]. In our model, Nrf2 activation by sulforaphane treatment promoted the hybrid E/M phenotype and enhanced proliferation (Supplementary Figure S7). Additional signaling pathways and molecular programs, such as stemness and metabolic switching [48–50], may also be involved in the regulation of the cancer invasion process. All these factors can contribute to the migratory behavior in a context specific manner, and the interrelated roles of Nrf2 on EMT and Notch signaling may explain the discrepancy on the functions of Nrf2 on the collective cancer migration.

This study applied an integrated experimental-computation approach to investigate the function of Nrf2 in collective cancer migration. The computation prediction based on our theoretical frameworks generally captured the observed hybrid E/M phenotypes. We note several limitations of the study. For instance, the current model could reproduce well the response of EMT and Notch signaling upon Nrf2 upregulation via sulforaphane, but could not capture the decrease of Dll4 and increase of Notch1 observed in the Nrf2-KO, potentially pointing to loss of adhesion and weakening of Notch signaling between mesenchymal migrating cells as an important element to integrate into future modeling efforts. The complex interplay between signaling and migratory dynamics also underscores future theoretical challenges that are not explicitly considered in our current model, including (1) the coupling of biochemical and mechanical regulation of cell migration, (2) the effect of cell proliferation on cell patterning, and (3) the context-specificity of the EMT program in terms of both transcriptional response and number of intermediate phenotypes in the EMT spectrum [51–53]. Furthermore, in order to overcome the limitations of using pharmacological methods such as sulforaphane, additional experimental models involving specific ways of perturbing one or multiple genes (i.e., gene editing techniques such as CRIPSR/Cas9), epistasis studies modulating Notch signaling and the EMT network, and more physiologically relevant invasion models should be performed to investigate the impact of the Nrf2-EMT-Notch network on collective cancer invasion. Future experimental and computational investigations will be required to fully understand the role of Nrf2 on cancer invasion.

## Supplementary Materials

The following are available online at www.mdpi.com/xxx/s1, Mathematical model of EMT-Nrf2-Notch signaling. Table 1. Nucleic sequences for the dsLNA biosensor. Table 2. Parameters of the Nrf2-EMT circuit. Table 3. Parameters of the Notch circuit. Table 4. Newly introduced parameters for NICD-Nrf2-Notch feedback. Table 5. Parameters for translation, mRNA degradation and microRNA degradation upon protein-microRNA binding. Figure S1. Double-stranded locked nucleic acid biosensor schematic. Figure S2. The intensity product of E-cadherin and ZEB1 near the leading edge. Figure S3. Spatiotemporal patterning as a function of external EMT induction. Figure S4. Spatial distribution of EMT phenotypes as a function of Notch signaling strength. Figure S5. Analysis of leading edge in the lattice model. Figure S6. Notch signaling in the lattice model. Figure S7. NRF2 modulates cell proliferation near the wound edge. Movies 1-3. Spatiotemporal dynamics of the lattice model for medium, low, and high levels of Nrf2 induction.

## Author Contributions

“Conceptualization, H.L., M.J., and P.W.; software, F.B., H.L., J.O., M.J.; investigation, S.M., N.Z., and F.B.; writing—original draft preparation, S.M., F.B., P.W.; writing—review and editing, N.Z., H.L., J.O., and M.J.; All authors have read and agreed to the published version of the manuscript.”

## Data Availability Statement

All the data supporting the findings of this study are available within the article and its supplementary information files and from the corresponding author upon reasonable request.

