## Supplementary Information for "Nrf2 modulates the hybrid epithelial/mesenchymal phenotype and Notch signaling during collective cancer migration"

### A. Mathematical model of EMT-Nrf2-Notch signaling

1) Intracellular EMT-Nrf2 signaling

We have previously introduced a mathematical model to describe the intracellular signaling dynamics of the networks controlling EMT and Nrf2[1]. The coupled dynamics of all species in the circuit (miR-34: W, miR-200: Y, Snail: S, ZEB1: Z, Keap1: K, E-cadherin: E, Nrf2: X) is described by coupled ordinary differential equations:

$$\frac{d\mu_{34}}{dt} = g_{\mu_{34}} H^{-}(S) H^{-}(Z) - g_S H^S(S) H^S(I) H^S(I_{ext}) P_y(\mu_{34}, 2) - \gamma_{\mu_{34}} \mu_{34} \quad (1a)$$
$$\frac{d\mu_{200}}{dt} = g_{\mu_{200}} H^{-}(Z) H^{-}(S) - g_Z H^S(Z) H^S(S) H^{-}(E) P_y(\mu_{200}, 6) - \gamma_{\mu_{200}} \mu_{200} \quad (1b)$$
$$\frac{dZ}{dt} = k_P g_Z H^{+}(Z) H^{+}(S) H^{-}(E) P_l(\mu_{200}, 6) - \gamma_Z Z \quad (1c)$$
$$\frac{dS}{dt} = k_P g_S H^{-}(X) H^{-}(S) H^{+}(I_{ext}) P_l(\mu_{34}, 2) - \gamma_S S \quad (1d)$$
$$\frac{dK}{dt} = k_K H^{-}(\mu_{200}) - \gamma_K K \quad (1e)$$
$$\frac{dE}{dt} = k_E H^{-}(Z) - \gamma_E E \quad (1f)$$
$$\frac{dX}{dt} = k_X H^{-}(K) H^{-}(E) - \gamma_X X \quad (1g)$$

In eqs. (1a-d), $g_{\mu_{200}}$, $g_{\mu_{34}}$, $g_Z$and $g_S$ represent basal transcription rate constants for miR-200, miR-34, ZEB1 and Snail, while $k_P$ is the translation rate for ZEB1 and Snail. Since mRNA dynamics of Keap1, E-cadherin and Nrf2 are not considered explicitly, their transcription and translation rate constants are merged into a single parameter ($k_K, k_E, k_X$, respectively). Moreover, each species ($i$) degrades with its own degradation rate ($\gamma_i$).

Additionally, several transcriptional interactions are described by shifted Hill functions that multiply the basal production rates. In particular, following the circuit of Figure 2A: miR-34 and miR-200 are both inhibited by ZEB1 and Snail; ZEB1 self-activates, and is further activated by Snail and inhibited by E-cadherin; Snail can be induced by external EMT-inducers such as TGF-beta or Wnt (indicated by $I_{ext}$), self-inhibits its own transcription, and is inhibited by Nrf2; Nrf2 is inhibited by E-cadherin and Keap1.

Moreover, the microRNAs miR-34 and miR-200 inhibit Snail and ZEB1, respectively, at the post-translational level via noncoding RNA-mRNA binding and degradation. These interactions further decrease the levels of miR-34 and miR-200 due to degradation of the microRNA-mRNA complex. Specifically, the production rates of ZEB1 and Snail are weighted by modulating functions $P_l(\mu_{200}, 6)$, $P_l(\mu_{34}, 2)$, whereas the equations for miR-34 and miR-200 include loss terms proportional to ZEB1, Snail production rates and modulated by functions $P_y(\mu_{34}, 2)$, $P_y(\mu_{200}, 6)$. The explicit form and derivation of these functions is discussed in the following section A4.

All parameters for the Nrf2-EMT circuit were taken from our previous modeling [1] and are summarized in Table 2.

2) Cell-cell communication via Notch signaling

To model cell-cell signaling through Notch, we start from the modeling framework developed by Boareto and collaborators [3,4]. The model is general and refers to the behavior of different Notch, Delta and Jagged subtypes. In the main text, we substitute Notch, Delta and Jagged with the specific subtypes that were probed experimentally: Notch1, Dll4 and Jag1. In this model, the molecular copy numbers of Notch (N), Delta (D), Jagged (J), and Notch Intracellular domain (I or NICD) are described by a system of ODEs:

$$\frac{dN}{dt} = k_P g_N H^{S+}(I) - N[(k_c D + k_c J) + [(k_t D_{ext} + (k_t J_{ext})] - \gamma N \quad (2a)$$

$$\frac{dD}{dt} = k_P g_D H^{S-}(I) - D\,(k_c N + k_t N_{ext}) \; - \gamma D \quad (2b)$$

$$\frac{dJ}{dt} = k_P g_J H^{S+}(I) - J\,(k_c N + k_t N_{ext}) - \gamma J \quad (2c)$$

$$\frac{dI}{dt} = N(k_t D_{ext} + k_t J_{ext}) - \gamma_I I \quad (2d)$$

Notch receptors can bind to external Delta and Jagged ligands ($D_{ext}, J_{ext}$) with transactivation rate $k_t$. These external ligands can represent soluble molecules in the cellular microenvironment as well as transmembrane ligands at the surface of neighboring cells. Similarly, Delta and Jagged transmembrane ligands can bind to external Notch receptors ($N_{ext}$). Additionally, the model considers binding between ligands and receptors within the same cell with rate constant $k_c$, which does not lead to downstream regulation but only degradation of the ligand-receptor complex (typically referred to as "cis-inhibition"). Transactivation of the Notch receptor leads to release of NICD, which in turns transcriptionally activates Notch and Jagged while inhibiting Delta (modeled via Hill functions). Finally, Notch, Delta, and Jagged degrade with rate constant $\gamma$, while NICD degrades with a faster rate $\gamma_I$. All parameters for the Notch circuit were taken from previous publications [3,4] and are summarized in Table 3.

3) Integrated EMT-Nrf2-Notch circuit

We integrate the Nrf2-EMT and Notch circuit as follows. The coupling between Notch and EMT circuits has been previously modeled by Boareto and collaborators [2], and includes the following additional interactions (also shown in Figure 2B).

- NICD transcriptionally activates Snail, thus resulting in a shifted Hill function multiplying the production rate of Snail ($H^{+}(I)$).

- miR-34 inhibits Notch and Dll, while miR-200 inhibits Jag. Similar to the miR-34/Snail and miR-200/ZEB1 interactions, these connections are modeled with the microRNA-TF chimera formalism, thus resulting in additional functions $P_l, P_y$.

Moreover, we introduce a specific interaction between the Notch and Nrf2 modules based on information from literature. NICD inhibits Nrf2, which in turns inhibits Notch, thus resulting in shifted Hill functions that modulate the production rate of Nrf2 ($H^-(I)$) and Notch ($H^-(X)$), respectively.

The full set of equation is (newly introduced terms are marked in red):

Nrf2-EMT circuit:

$$\frac{d\mu_{34}}{dt} = g_{\mu_{34}} H^-(S)H^-(Z) - g_S H^S(S)H^S(I)H^S(I_{ext})\, H^{S+}(I) P_y(\mu_{34}, 2)$$
$$-k_P g_N H^{S+}(I)H^{S-}(X)P_y(\mu_{34}, 2) - k_P g_D H^{S-}(I)P_y(\mu_{34}, 3) - \gamma_{\mu_{34}}\mu_{34} \quad (3a)$$

$$\frac{d\mu_{200}}{dt} = g_{\mu_{200}} H^-(Z)H^-(S) - g_Z H^S(Z)H^S(S)H^-(E)P_y(\mu_{200}, 6)$$
$$-k_P g_J H^{S+}(I)P_l(\mu_{200}, 5) - \gamma_{\mu_{200}}\mu_{200} \quad (3b)$$

$$\frac{dZ}{dt} = k_P g_Z H^+(Z)H^+(S)H^-(E)P_l(\mu_{200}, 6) - \gamma_Z Z \quad (3c)$$

$$\frac{dS}{dt} = k_P g_S\, H^-(X)\, H^-(S)\, H^+(I_{ext})\, H^{S+}(I)\, P_l(\mu_{34}, 2) - \gamma_S S \quad (3d)$$

$$\frac{dK}{dt} = k_K H^-(\mu_{200}) - \gamma_K K \quad (3e)$$

$$\frac{dE}{dt} = k_E H^-(Z) - \gamma_E E \quad (3f)$$

$$\frac{dX}{dt} = k_X\, H^-(K)\, H^-(E)\, H^{S-}(I) - \gamma_X X \quad (3g)$$

Notch circuit:

$$\frac{dN}{dt} = k_P g_N H^{S+}(I)H^{S-}(X)P_l(\mu_{34}, 2) - N[(k_c D + k_c J) + [(k_t D_{ext} + (k_t J_{ext})] - \gamma N \quad (3h)$$

$$\frac{dD}{dt} = k_P g_D H^{S-}(I)P_l(\mu_{34}, 3) - D\,(k_c N + k_t N_{ext}) - \gamma D \quad (3i)$$

$$\frac{dJ}{dt} = k_P g_J H^{S+}(I)P_l(\mu_{200}, 5) - J\,(k_c N + k_t N_{ext}) - \gamma J \quad (3j)$$

$$\frac{dI}{dt} = N(k_t D_{ext} + k_t J_{ext}) - \gamma_I I \quad (3k)$$

The newly introduced parameters include the double negative feedback between NICD, Nrf2 and Notch, and are summarized in Table 4.

4) Modeling of microRNA mediated inhibition

Post-translational inhibition by microRNAs (miR-200, miR-34) is described using the MBC framework (microRNA-Based-Chimeric circuits) first introduced by Lu and collaborators[5] with the abovementioned functions $P_l(\mu, n)$ and $P_y(\mu, n)$. The function $P_l(\mu, n)$ models post-translational inhibition exerted by a microRNA species $\mu$ on a target species. Here, $n$ is the number of binding sites for microRNA molecules on the target species. Similarly, $P_y(\mu, n)$ describes the microRNA decrease due to microRNA/mRNA complex degradation:

$$P_l(\mu, n) = \frac{L(\mu, n)}{Y_m(\mu, n) + k_m} \quad (4a)$$

$$P_y(\mu, n) = \frac{Y_\mu(\mu, n)}{Y_m(\mu, n) + k_m} \quad (4b)$$

where $L(\mu, n)$ is the total translation rate, $Y_m(\mu, n)$ is the total active degradation rate of target mRNA and $Y_\mu(\mu, n)$ is the total degradation rate of the microRNA:

$$L(\mu, n) = \sum_{i=0}^{n} l_i C_i{}^n M_i{}^n(\mu) \quad (5)$$

$$Y_m(\mu, n) = \sum_{i=0}^{n} \gamma_{mi} C_i{}^n M_i{}^n(\mu) \quad (6)$$

$$Y_\mu(\mu, n) = \sum_{i=0}^{n} \gamma_{\mu i} C_i{}^n M_i{}^n(\mu) \quad (7)$$

In eqs. (5-7), $l_i$, $\gamma_{mi}$ and $\gamma_{\mu i}$ are individual rates for the case of $i$ microRNAs bound to the protein. Also, $C_i{}^n$ is the number of arrangements of $i$ molecules of microRNA in $n$ binding sites:

$$C_i{}^n = \frac{n!}{i!\,(n-i)!} \quad (8)$$

and

$$M_i{}^n(\mu) = \frac{\left(\frac{\mu}{\mu_0}\right)^i}{\left(1 + \frac{\mu}{\mu_0}\right)^n} \quad (9)$$

All rate parameters for MBC modelling are summarized in table 5.

5) Simulation details and identification of EMT phenotypes

The 150x50 hexagonal lattice is prepared with randomized initial conditions. The values of Notch1, Dll4, Jagged1, NICD, miR-34, miR-200, ZEB1, Snail, E-cad, Keap1, and Nrf2 are sampled from a uniform distribution bound between 0 and a maximal value that depends on the variable. The maximal values are estimated based on the largest possible steady state level that each variable can assume based on the model's parameters (i.e., in a case where all inhibitory interactions are neglected and all positive interactions are maximized). At t=0 the EMT-inducer is fixed at a constant level ($I_{EXT}$) at the leftmost edge of the lattice and is equal to zero everywhere else in the lattice.

Starting from the randomized initial condition, the ODEs describing each cell and the reaction-diffusion equation for the EMT-inducer are integrated simultaneously with a forward Euler scheme. The model has periodic boundary conditions along the y-axis (i.e., the top and bottom rows of cells are in contact); conversely, the leftmost and rightmost boundaries are free to model the spatial organization of the cell layer.

Based on previous analysis of steady state in the EMT circuit, we classify cells as epithelial (E), hybrid E/M or mesenchymal (M) based on the level of miR-200 (E: miR-200>15000 molecules; E/M: 5000 molecules<miR-200<15000 molecules; M: miR-200<5000 molecules). These thresholds were derived by studying the bifurcation diagram of a single cell version of the model. In this single cell bifurcation diagram, an 'epithelial' branch always satisfied miR-200>15000; the hybrid E/M branch always satisfied 15000<miR-200<5000; and the mesenchymal branch always satisfied miR-200<5000. Further information can be found in the original model by Lu and collaborators[5].

### B. Supplementary Tables

Table 1. Nucleic sequences for the dsLNA biosensor*

| | |
|---|---|
| $\beta$-actin Probe | /5Alex647N/+AG+GT+TT+TG+TC+AA+GA+AA+GG+GT |
| $\beta$-actin Quencher | GACAAAACCT/3IAbRQSp/ |
| Random Probe | /5Alex647N/+AC+TC+CA+CT+TG+TT+CA+CC+GA+TA |
| Random Quencher | CAAGTGGAGT/3IAbRQSp/ |
| miR-200c-3p Probe | /5Alex647N/+TC+CA+TC+AT+TA+CC+CG+GC+AG+TA+TT+A |
| miR-200c-3p Quencher | GGTAATGATGGA/3IAbRQSp/ |
| Notch1 Probe | /5Alex647N/+TG+CG+GT+CT+GT+CT+GG+TT+GT+GC |
| Notch1 Quencher | ACAGACCGCA/3IAbRQSp/ |
| Dll4 Probe | /5Alex647N/+AA+GG+GC+AG+TT+GG+AG+AG+GG+TT |
| Dll4 Quencher | AACTGCCCTT/3IAbRQSp/ |

* Sequences are in the 5’ to 3’ direction; + indicates LNA monomers

Table 2. Parameters of the Nrf2-EMT circuit. (*) Production rates for Keap1, E-cadherin and Nrf2 include both transcription and translation and are therefore much larger.

| Parameter group | Parameter | Value | Dimensions |
|---|---|---|---|
| Degradation | $\gamma_{\mu200}, \gamma_{\mu34}, \gamma_Z, \gamma_S, \gamma_K, \gamma_E, \gamma_X$ | $0.05, 0.05, 0.1, 0.1, 0.1, 0.1, 0.1$ | $h^{-1}$ |
| Production | $k_P, g_{\mu200}, g_{\mu34}, g_Z,$ $g_S, g_K, g_E, g_X$ | $100, 2100, 1350, 11, 90,$ $50000, 50000, 50000$ (*) | $h^{-1}$ |
| Hill coefficient | $n_{S,\mu34}, n_{Z,\mu34}, n_{S,\mu200}, n_{Z,\mu200},$ $n_{Z,Z}, n_{S,Z}, n_{S,S}, n_{X,S}, n_{I,S},$ $n_{\mu200,K}, n_{Z,E}, n_{K,X}, n_{E,X}$ | $1, 3, 2, 3,$ $2, 2, 1, 2, 2,$ $2, 2, 2, 2$ | $Dimensionless$ |
| Hill fold-change | $\lambda_{S,\mu34}, \lambda_{Z,\mu34}, \lambda_{S,\mu200}, \lambda_{Z,\mu200},$ $\lambda_{Z,Z}, \lambda_{S,Z}, \lambda_{S,S}, \lambda_{X,S}, \lambda_{I,S},$ $\lambda_{\mu200,K}, \lambda_{Z,E}, \lambda_{K,X}, \lambda_{E,X}$ | $0.1, 0.2, 0.1, 0.1,$ $7.5, 10, 0.1, 0.67, 10,$ $0.1, 0.1, 0.33, 0.33$ | $Dimensionless$ |
| Hill threshold | $I_{S,\mu34}, I_{Z,\mu34}, I_{S,\mu200}, I_{Z,\mu200},$ $I_{Z,Z}, I_{S,Z}, I_{S,S}, I_{X,S}, I_{I,S},$ $I_{\mu200,K}, I_{Z,E}, I_{K,X}, I_{E,X}$ | $300K, 600K, 180K, 220K,$ $25K, 180K, 200K, 1000K, 50K,$ $5K, 20K, 250K, 250K$ | $Molecules$ |

Table 3. Parameters of the Notch circuit.

| Parameter group | Parameter | Value | Dimensions |
|---|---|---|---|
| Degradation | $\gamma_N, \gamma_D, \gamma_J, \gamma_I$ | $0.1, 0.1, 0.1, 0.5$ | $h^{-1}$ |
| Production | $k_P, g_N, g_D, g_J$ | $100, 8, 20, 70$ | $h^{-1}$ |
| Hill coefficient | $n_{I,N}, n_{I,D}, n_{I,J}$ | $2, 2, 5, 2, 2$ | $Dimensionless$ |
| Hill fold-change | $\lambda_{I,N}, \lambda_{I,D}, \lambda_{I,J}$ | $2, 0, 2$ | $Dimensionless$ |
| Hill threshold | $I_{0,N}, I_{0,D}, I_{0,J}$ | $200, 200, 200$ | $Molecules$ |
| Binding rates | $k_T, k_C$ | $10^{-5}, 10^{-4}$ | $h^{-1}Molecules^{-1}$ |
| Fringe activity | $n_F, \lambda_{F,D}, \lambda_{F,J}$ | $1.0, 3.0, 0.3$ | $Dimensionless$ |

Table 4. Newly introduced parameters for NICD-Nrf2-Notch feedback.

| Parameter group | Parameter | Value | Dimensions |
|---|---|---|---|
| Hill coefficient | $n_{X,N}, n_{I,X}$ | $2, 2$ | $Dimensionless$ |
| Hill fold-change | $\lambda_{X,N}, \lambda_{I,X}$ | $2.5, 2.5,$ | $Dimensionless$ |
| Hill threshold | $I_{X,N}, I_{I,X}$ | $750K, 200$ | $Molecules$ |

Table 5. Parameters for translation, mRNA degradation and microRNA degradation upon protein-microRNA binding.

| **Parameter group** | **Parameter** | **Value** | **Dimensions** |
|---|---|---|---|
| Translation rate | $l_i$ | $1.0, 0.6, 0.3, 0.1, 0.05, 0.05, 0.05$ | |
| mRNA degradation rate | $\gamma_{mi}$ | $0, 0.04, 0.2, 1.0, 1.0, 1.0, 1.0$ | $h^{-1}$ |
| microRNA degradation rate | $\gamma_{\mu i}$ | $0, 0.005, 0.05, 0.5, 0.5, 0.5, 0.5$ | $h^{-1}$ |

### C. Supplementary figures

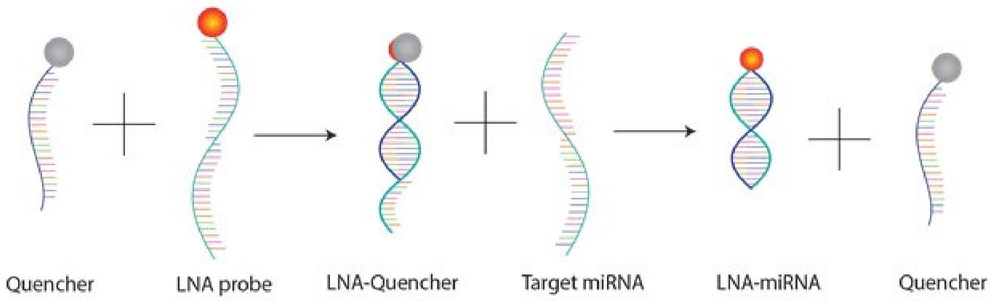


**Figure S1.** Double-stranded locked nucleic acid biosensor schematic. Illustration of the working principle for the double-stranded locked nucleic acid (dsLNA) biosensor. The quencher-labeled LNA sequence is hybridized with the fluorophore-labeled LNA probe to form the dsLNA biosensor. The homogeneous biosensor is transfected into cancer cells. With the presence of a target mRNA or miRNA, the quencher probe displaced by the target, allowing the fluorophore-labeled LNA probe to fluoresce.

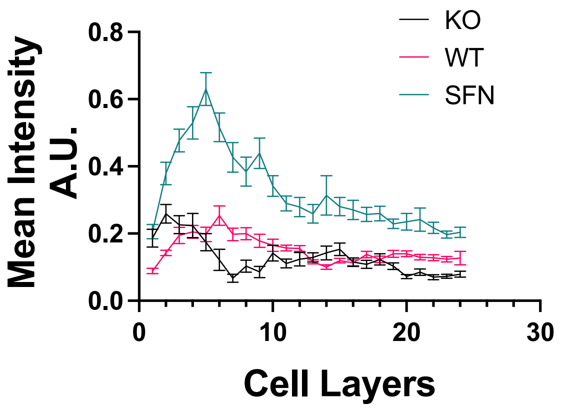


**Figure S2.** The intensity product of E-cadherin and ZEB1 near the leading edge. To analyze the hybrid E/M cells, the intensity product of E-cadherin and ZEB1 (E*Z) was calculated. The average value was enhanced with sulforaphane treatment (SFN) and reduced with Nrf2 KO.

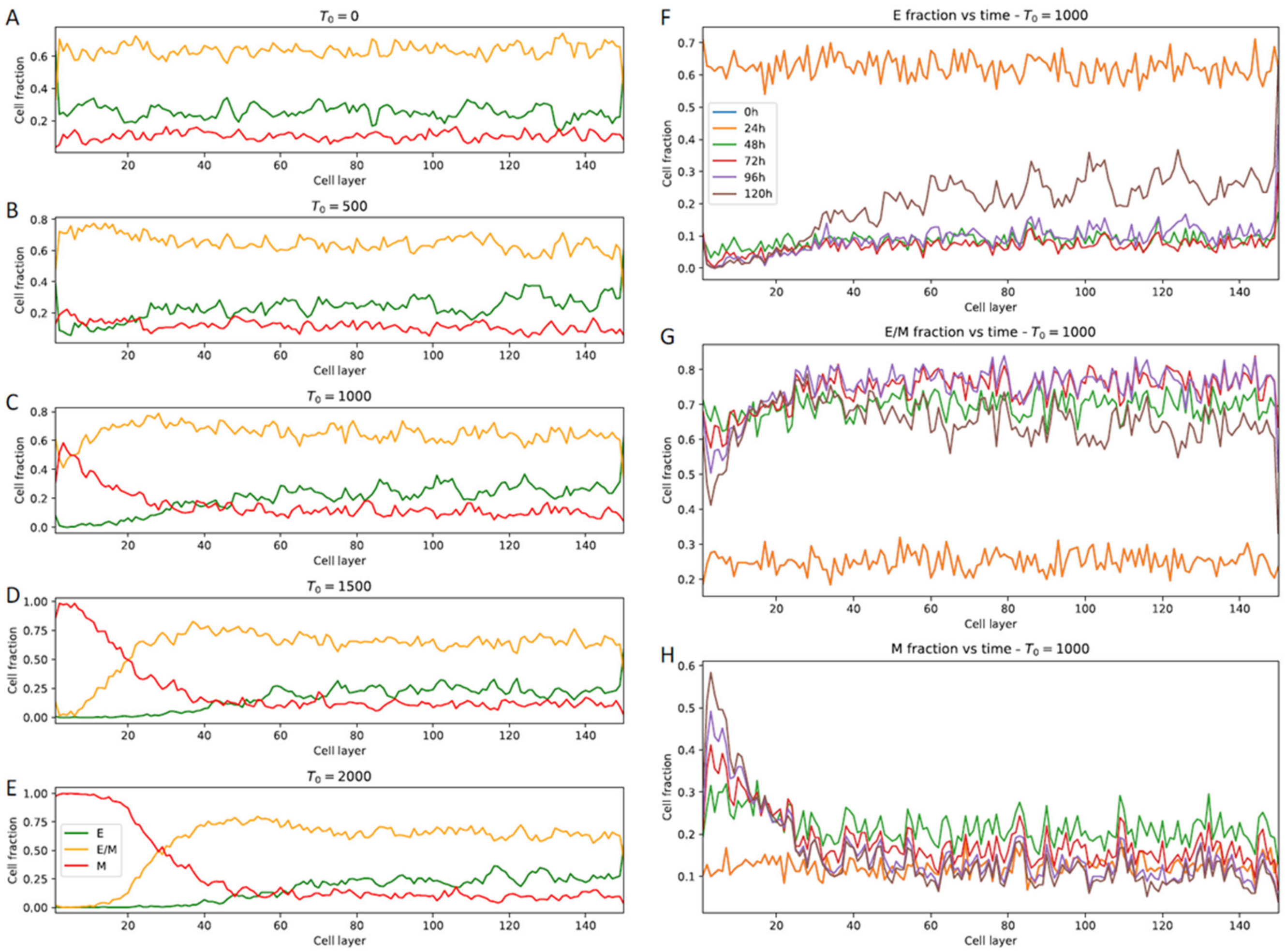


**Figure S3.** Spatiotemporal patterning as a function of external EMT induction. (A-E) Fraction of E, hybrid E/M, and M cells as a function of distance from the invasive edge for increasing levels of external EMT inducer ($T_0$). $T_0$ represents the fixed level of EMT inducer at the leftmost end of the lattice (the invasive edge), which then diffuses throughout the lattice. These fractions are computed after 120 hours of simulation starting from randomized initial conditions. (F-H) Fraction of E, hybrid E/M, and M cells as a function of distance from the invasive edge at different time points over the course of the simulation. All results are averaged over 5 simulations starting from randomized initial conditions.

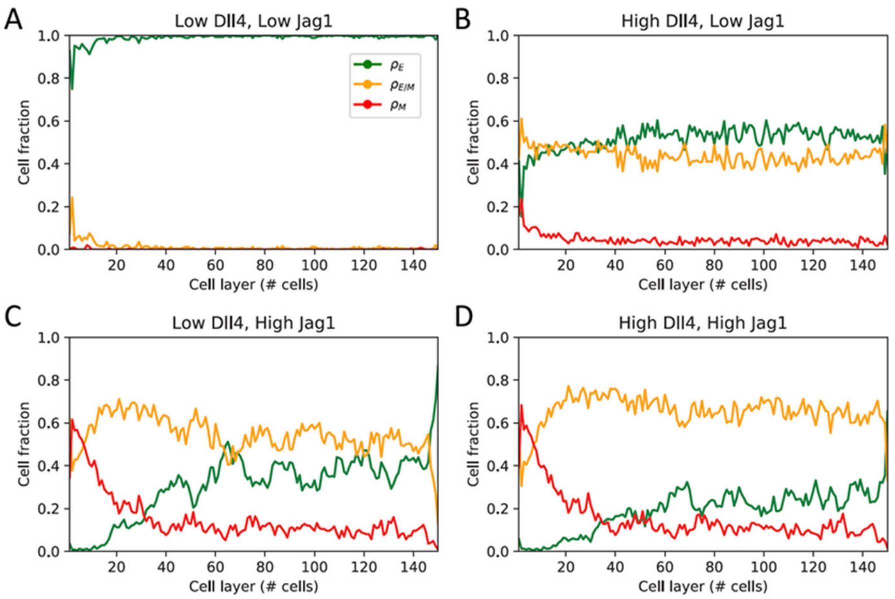


**Figure S4.** Spatial distribution of EMT phenotypes as a function of Notch signaling strength. (A) Fraction of E, E/M, and M cells as a function of distance from the invading edge for a case where both Notch1-Dll4 signaling and Notch1-Jagged1 signaling are weak ($g_D = 20\ molec/h$, $g_J = 20\ molec/h$). (B) Same as (A) for strong Notch1-Dll4, weak Notch1-Jagged1 ($g_D = 120\ molec/h$, $g_J = 20\ molec/h$). (C) Same as (A) for weak Notch1-Dll4, strong Notch1-Jagged1 ($g_D = 20\ molec/h$, $g_J = 60\ molec/h$). (D) Same as (A) for strong Notch1-Dll4, strong Notch1-Jagged1 ($g_D = 120\ molec/h$, $g_J = 60\ molec/h$). For all cases, the external EMT inducing signal and Nrf2 production rate are at a medium level ($T_0 = 1000\ molec$, $g_{Nrf2} = 0.5\ \times 10^5\ molec/h$). All results are averaged over 5 simulations starting from randomized initial conditions.

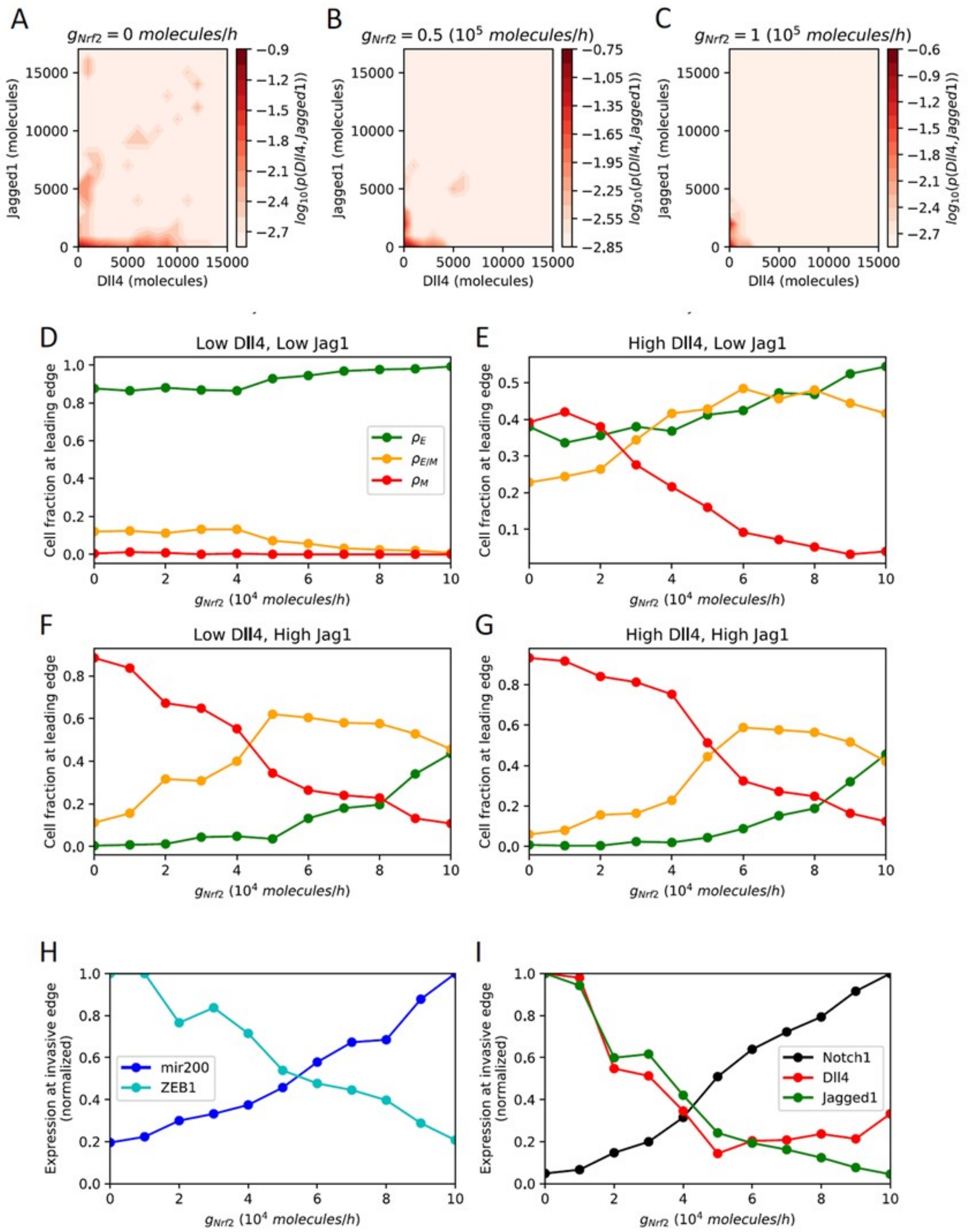


**Figure S5.** Analysis of leading edge in the lattice model. (A-C) Log-normalized probability to observe cells with varying levels of Dll4 and Jag4 in the invasive edge under weak, medium, and strong Nrf2 induction. (D-G) Fractions of E, hybrid E/M and M cells in the leading edge as a function of Nrf2 production rate. The four panels show different combinations of weak/strong Notch1-Dll4 and Notch1-Jagged1 signaling (same parameter combinations of Figure S4). (H) Average expression of miR-200 and ZEB1 at the leading edge as a function of Nrf2 production rate for a case of low Dll4, high Jagged1 signaling. (I) Average expression of Notch1, Dll4 and Jagged1 at the leading edge as a function of Nrf2 production rate. All results are averaged over 5 simulations starting from randomized initial conditions.

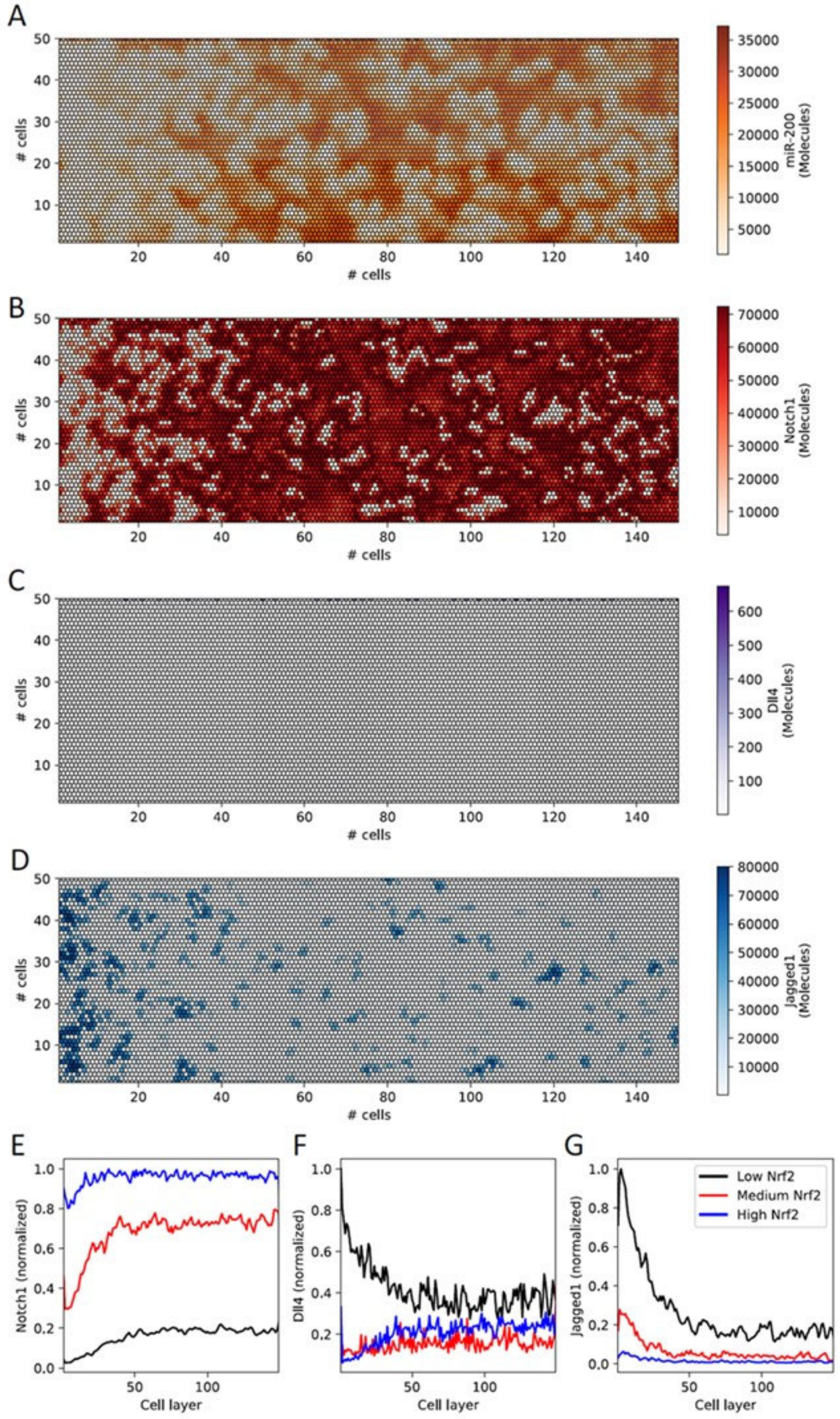


**Figure S6.** Notch signaling in the lattice model. (A-D) Heatmap showing the multicell patterns of miR-200, Notch1, Dll4 and Jagged1 for a medium Nrf2 activation ($g_{Nrf2} = 0.5 \times 10^5\, molec/h$). (E-G) Average expression of Notch1, Dll4 and Jagged1 as a function of distance from the invading edge for low, medium and high Nrf2 induction ($g_{Nrf2} = 0\, molec/h$, $g_{Nrf2} = 0.5 \times 10^5\, molec/h$, $g_{Nrf2} = 10^5\, molec/h$, respectively). Results for (E-G) are averaged over 5 simulations starting from randomized initial conditions.

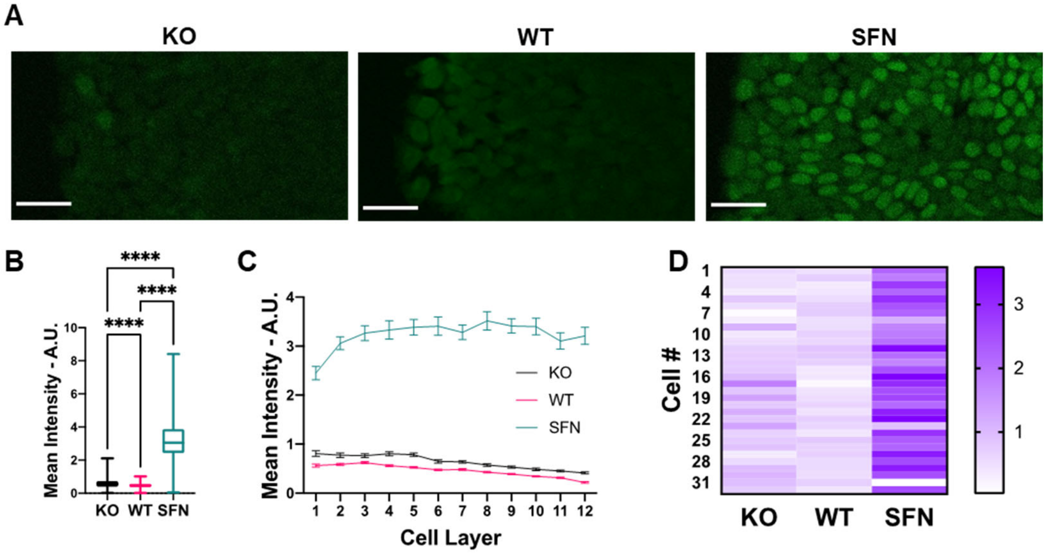


**Figure S7.** Nrf2 modulates cell proliferation near the wound edge. (A) Cell proliferation assay for CRISPR/Cas9 NFE2L2-KO Pool RT4 cells (KO), wild-type RT4 cells (WT) and sulforaphane treated RT4 cells (SFN), respectively. Scale bars, 50 µm. (B-D) Quantification of proliferation data. (B) Average intensity over the whole cell layer for KO, WT, and SFN, respectively. (C) Tracing of relative fluorescence intensity for KO, WT, and SFN per tissue depth measured as number of cell layers. (D) Heatmap of representative cells in the first row for KO, WT, and SFN, respectively. The nonparametric Kruskal-Wallis test along with the Dunn's multiple comparisons test were used to compare across groups (view Materials and Methods section). For each experiment n > 500 cells per condition.

### D. Supplementary movies

M1: Spatiotemporal dynamics of the lattice model for a medium level of Nrf2 induction ($g_{Nrf2} = 0.5 \times 10^5\, molec/h$). Top: external EMT-inducing signal as a function of distance from the leading edge. Middle: Spatial patterning of epithelial (green), hybrid E/M (yellow) and mesenchymal (red) cells. Bottom: Fraction of epithelial (green), hybrid E/M (yellow) and mesenchymal (red) cells as a function of distance from the leading edge.

M2: Spatiotemporal dynamics of the lattice model for a low level of Nrf2 induction ($g_{Nrf2} = 0\, molec/h$). Top: external EMT-inducing signal as a function of distance from the leading edge. Middle: Spatial patterning of epithelial (green), hybrid E/M (yellow) and mesenchymal (red) cells. Bottom: Fraction of epithelial (green), hybrid E/M (yellow) and mesenchymal (red) cells as a function of distance from the leading edge.

M3: Spatiotemporal dynamics of the lattice model for a high level of Nrf2 induction ($g_{Nrf2} = 10^5\, molec/h$). Top: external EMT-inducing signal as a function of distance from the leading edge. Middle: Spatial patterning of epithelial (green), hybrid E/M (yellow) and mesenchymal (red) cells. Bottom: Fraction of epithelial (green), hybrid E/M (yellow) and mesenchymal (red) cells as a function of distance from the leading edge.
